# ORC binding to Heterochromatin Protein 1 through intrinsically disordered regions is required for heterochromatin structure and function

**DOI:** 10.64898/2026.07.17.738698

**Authors:** Yuchang Luo, Srivarsha Rajshekar, Harish Kumar, James M. Berger, Gary H. Karpen, Michael R. Botchan

## Abstract

The Origin Recognition Complex (ORC) is known for initiating DNA replication in eukaryotic cells, but increasing evidence suggests that ORC has multiple functions. Here we show that the organization and function of two major nuclear components, heterochromatin and the nucleolus, depend on multivalent interactions between Orc1 and Heterochromatin Protein 1a (HP1a) in *Drosophila melanogaster*. Specifically, binding requires two short motifs (R1 and R2) in the intrinsically disordered region (IDR) of Orc1, and two motifs (HGM and CTE) located in the N- and C-terminal regions of HP1a. Pairing of these four motifs promotes ORC–HP1 interactions, which also requires HP1 dimerization. Disrupting ORC–HP1a interactions by mutating the R1/R2 motifs causes defects in heterochromatin functions, specifically suppression of Position-Effect Variegation (PEV), ribosomal DNA (rDNA) decondensation, increased rRNA transcription, and nucleolar expansion, without global loss of H3K9me2 epigenetic modifications. Our findings indicate that ORC acts as a structural regulator that organizes heterochromatic regions and promotes proper genome regulation through multivalent interactions with HP1 and chromatin.

## Introduction

Many essential cellular molecules and complexes regulate multiple, often unrelated functions. For example, the six-subunit Origin Recognition Complex (ORC), an essential enzyme in the AAA+ protein superfamily, is ubiquitous in eukaryotes and required before S-phase to recognize and license sites for DNA replication, by loading the latent MCM 2-7 complex ^1^. However, in metazoans, individual ORC subunits or the entire complex mediate functions outside of replication. For example, in humans and *Drosophila melanogaster* (*Dm*), cytoplasmic Orc6 subunit plays a direct role in cytokinesis through its interactions with a septin protein ^2,3^. In *Drosophila*, the Orc3 subunit is localized to synaptic buttons, and may regulate synaptic plasticity ^4,5^. Finally, ORC plays conserved roles in genome organization and gene regulation ^6^, and in mitotic chromosome condensation ^7,8^. These and other observations raise an important question: how does ORC mediate multiple interactions and functions within the cytoplasm and nucleus, independent from its role in replication?

Insights into mechanisms that mediate ORC’s non-replication functions in the nucleus came from biochemical, genetic and cytological studies in *Drosophila* ^6^. Pak et al. (1997) showed that *Dm* ORC co-localizes cytologically with heterochromatic regions of the genome in interphase and mitosis. Heterochromatin is a major component of most eukaryotic genomes with key roles in nuclear architecture, genome stability, transposon and gene silencing, and chromosome segregation. Heterochromatic regions are highly enriched for repetitive DNA (satellites, transposons), as well as nucleosomes containing H3K9me2/3 and its binding partner Heterochromatin Protein 1 (HP1, or HP1a in *Drosophila*). Biochemically, HP1a binds to the ORC complex through a domain within the largest subunit, Orc1 ^6^, which is now known as part of the intrinsically disordered region (IDR) of the protein ^9^. ORC–HP1 interactions are conserved in *Xenopus* ^6^ and humans ^10^. ORC haploinsufficiency, driven by an Orc2 null mutation, resulted in suppression of heterochromatin-mediated gene silencing (Position-Effect Variegation or PEV), suggesting that ORC, and specifically the Orc1 IDR, might impact heterochromatin structure and function through binding to HP1a. It is possible that ORC binding impacts heterochromatin by altering HP1-mediated compaction or other attributes of chromatin organization ^11^. Alternatively, both ORC ^9^ and HP1^12,13^ behave as biological membraneless condensates ^14,15^, formed by phase separation or similar mechanisms. Condensates are dynamic assemblies formed through multiple, weak interactions, and ORC binding could be required for HP1a condensate formation or biophysical properties, such as selective retention or exclusion of key regulators of heterochromatin functions^12^. However, many questions about ORC multifunctionality remain unanswered because the specific molecular basis for Orc1–HP1a binding are unknown, and we lack direct evidence that this interaction is critical for heterochromatin function *in vivo*.

To address these questions, here we used biochemical assays and gene editing techniques to precisely define ORC–HP1a binding mechanisms, and performed *in vivo* assessments of heterochromatin functions when Orc1–HP1a interactions were eliminated. The results show that Orc1–HP1a binding requires two short linear motifs (SLiMs) spaced closely within the *Dm*Orc1 IDR, and two short motifs located in the N- and C-terminal regions of HP1a. Biochemical experiments reveal that the Orc1 motifs bind the HP1a dimer through a non-canonical mechanism, distinct from previously described PxVxL-dependent HP1 Access Code (HAC) interactions with the HP1a dimer interface ^16^, formed by the chromoshadow domain and C-terminal extension. Importantly, disrupting Orc1–HP1a interactions by mutating these motifs at the endogenous *Drosophila orc1* locus results in suppression of PEV, as well as increased ribosomal DNA (rDNA) expression and nucleolar volume. Together, these results demonstrate that the Orc1 IDR SLiMs are required for HP1a binding, and in turn mediate important heterochromatin functions.

## Results

### Two separated SLiMs in Orc1 IDR mediate HP1a binding

Previous research demonstrated that *Drosophila* Orc1 binds HP1a through a 160-amino-acid region (K161 to S319, **Figure 1A**) ^6^, which starts at the end of the structured bromo-adjacent homology (BAH) domain, and extends into a long intrinsically disordered region (IDR, 187–549). To identify the molecular determinants within DmOrc1 that mediate its interaction with HP1a, we used a pull-down protocol where fragments of His_10_- (or His_10_-MBP-) tagged Orc1 proteins bound to beads were incubated with FLAG-tagged DmHP1a (**Supplementary Figure 1**, and **materials and methods**). Relative binding to WT in the pull-downs shows that Orc1 fragments containing residues 2–200 or 241–319 did not bind HP1a, whereas Orc1 (2–240) displayed robust HP1a binding (**Figure 1B**). Thus, deletion mapping demonstrates that Orc1 amino acids 200–240 contain critical recognition residues for its interaction with HP1a. Although such pull-down assays demonstrate interactions, it could not provide equilibrium measurements. We have used sensitive mass photometry ^17^ but could not measure a binding constant between DmOrc1 (2-240) and DmHP1a, consistent with weak and dynamic nature of IDR-mediated interactions.

**Figure 1.**
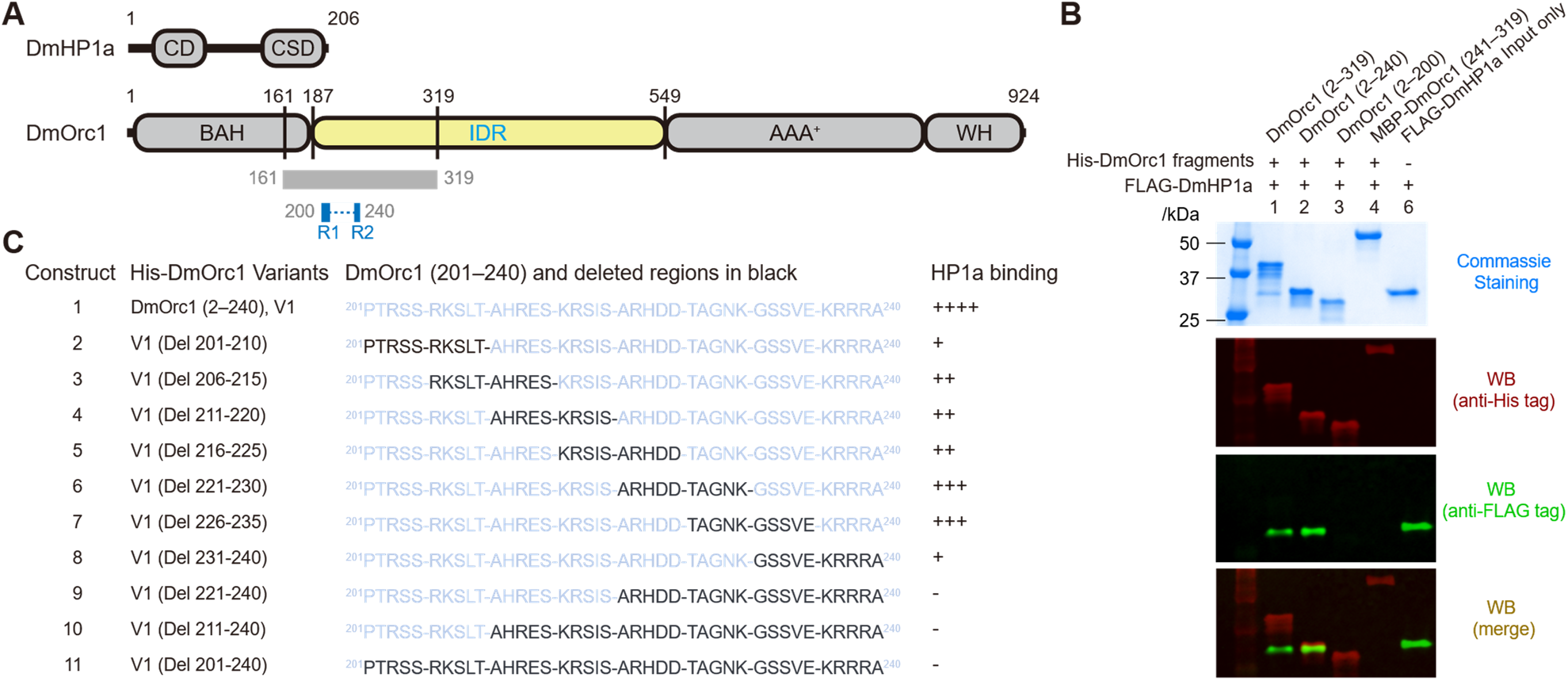
Identification of DmHP1a-binding regions within the DmOrc1 intrinsically disordered region. **(A)** Domain organization of *Drosophila melanogaster* HP1a (DmHP1a) and Orc1 (DmOrc1). DmHP1a: CD, chromodomain; CSD, chromoshadow domain; DmOrc1: BAH, bromo-adjacent homology domain; IDR, intrinsically disordered region; AAA^+^, ATPases associated with diverse activities; WH, winged-helix domain. The gray box indicates Orc1 residues 161–319, previously shown to mediate HP1a binding ^6^. The Orc1 Region 1 (R1, residues 200–209) and Regions 2 (R2, residues 234–239), HP1a-binding motifs defined in this study. **(B)** Binding of His_10_-tagged DmOrc1 fragments to FLAG-tagged DmHP1a. Coomassie-stained gel shows all the bound and released Orc1 and HP1a from pull-down assays. Immunoblotting detects His-tagged DmOrc1 fragments (red) and FLAG-tagged DmHP1a (green). **(C)** Deletion analysis of the Orc1 (2–240) identifies motifs within residues 201–210 and 231–240 that contribute to HP1a binding. Del denotes Delete; Black text indicates deleted regions. Qualitative relative binding of HP1a to wild-type Orc1 (2–240) is shown to the right; the relative signal intensities were categorized into five levels based on normalized values (wild-type set to 100%): –, <5%; +, 5–20%; ++, 20–40%; +++, 40–80%; and ++++, ≥80%. **Supplementary** Figure 2 shows the actual numerical quantitation for multiple measurements with error bars.

Next, to identify critical motifs within the 200–240 region of the Orc1 IDR, we generated a series of deletion mutations within His_10_-tagged DmOrc1 (2–240) fragment, including 10-aa deletions with 5-aa overlaps, and successively longer C-terminal deletions (**Figure 1C**). We quantified HP1a binding for the Orc1 deletion mutations relative to wild-type (WT) (See **Supplementary Figure 2)**, and classified impact on affinity as major (‘+’), modest (‘++’), or little to no effect (‘+++’ and ‘++++’) (**Figure 1C**). For example, deleting residues 201–210 or 231–240 caused a major reduction on Orc1–HP1a binding (+), whereas 211–220 caused a modest reduction (++) and deletions between 221–230 had little effect (+++) (**Figure 1C**, **Supplementary Figure 2**). We conclude that key binding determinants are concentrated near both ends of this 40-aa Orc1 IDR segment, mapping to residues 201–210 and 231–240, which correspond to two distinct 10-residue subregions. Furthermore, deletion of the internal segments of the Orc1 fragment (residues 216–235) showed robust binding to HP1 dimers (**Figure 1C**, **Supplementary Figure 2**) suggesting that the spacing between the two subregions is not critical, as might be expected for an intrinsically disordered region (IDR) given the flexible nature of IDR ^18^. In addition, based on the distribution of residues with bulky side-chain and the observation that deleting 226–235 had little effect on HP1a binding, the N-terminal determinant was expanded and defined as Region 1 (R1; residues 200–209), whereas the C-terminal determinant was refined to a shorter segment and defined as Region 2 (R2; residues 234–239) (**Figure 1A**).

To evaluate the role of R1 and R2 in HP1a binding, all bulky side-chain residues within R1 (R, P, K, and L) or R2 (V, E, K, and R) were simultaneously mutated, generating variants Mut R1 and Mut R2, respectively (**Figure 2A**). As the bulky side-chain residues in R1 are distributed throughout the sequence, they were substituted with alanine. In contrast, the six bulky side-chain residues in R2 form a contiguous cluster and were replaced with a combination of alanine and serine to reduce side-chain volume while avoiding the creation of an artificial poly-alanine tract. Both Mut R1 and Mut R2 exhibited markedly reduced HP1a binding, with Mut R1 displaying stronger impact. Importantly, combining these substitutions in a single mutant (Mut R1/R2) abolished detectable HP1a binding (**Figure 2A**). These results indicate that both R1 and R2 contribute to HP1a binding and act cooperatively to mediate the maximal interaction, with R1 making a larger contribution.

**Figure 2.**
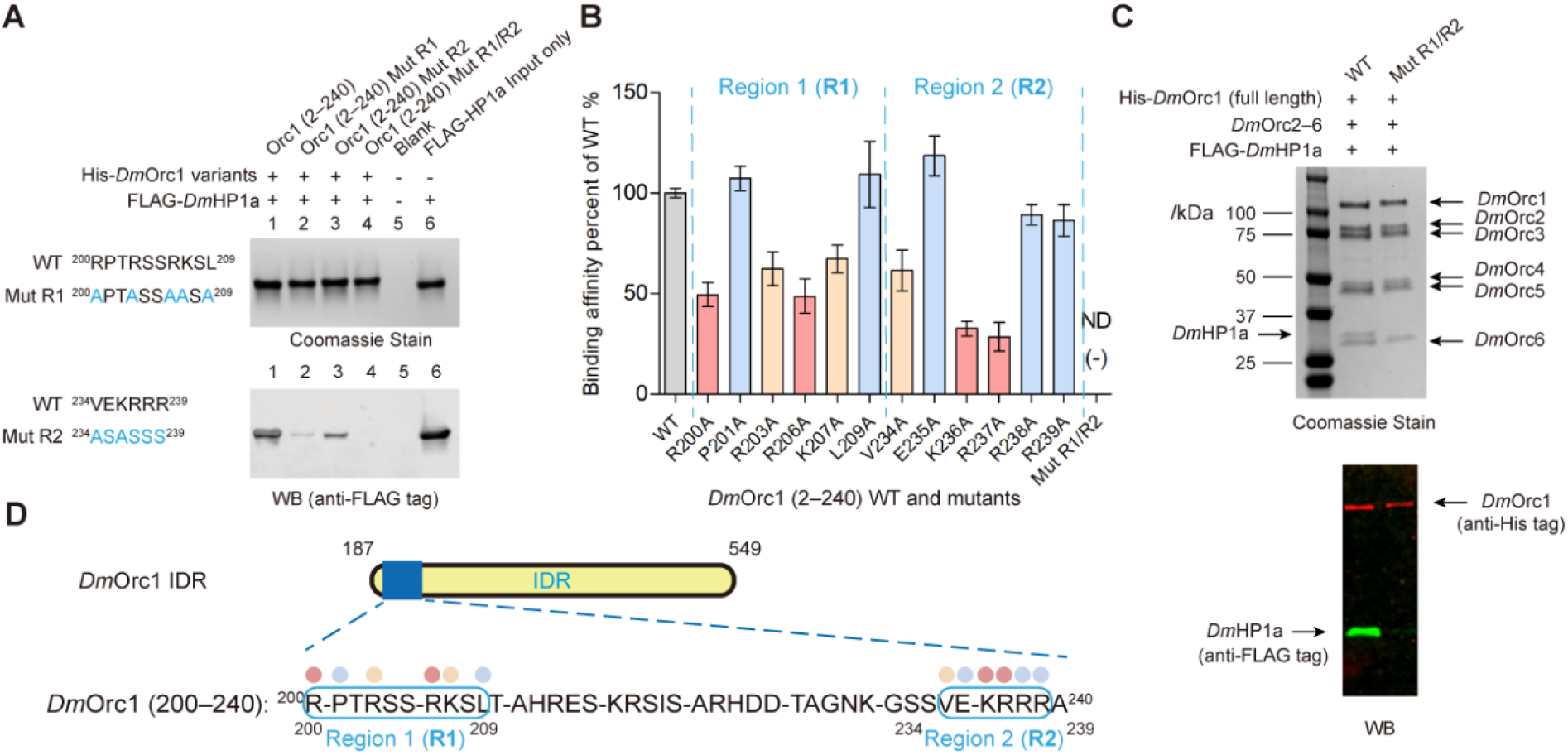
R1 and R2 motifs within the DmOrc1 intrinsically disordered region mediate binding to HP1a. **(A)** Mutating all large side-chain residues within R1 and/or R2 to small side-chain residues (alanine or serine) (Mut R1, Mut R2, Mut R1/R2) significantly reduces Orc1 (2–240) binding to HP1a, with Mut R1 having greater impact than Mut R2. No HP1a binding is detected when MutR1 and MutR2 are combined. Coomassie staining (top) and anti-FLAG immunoblotting to detect FLAG-HP1a (bottom) are shown. Mut R1 contains R200A, R203A, R206A, K207A, and L209A; Mut R2 contains V234A, E235S, K236A, R237S, R238S, and R239S. Mut R1/R2 represents a combination of the mutations in Mut R1 and Mut R2. **(B)** Quantification of binding for the indicated single alanine substitutions in Orc1 (2–240), displayed as a percentage of wild-type (WT) binding; error bars=standard error for 3 experiments. The combined Mut R1/R2 mutant shows no detectable binding (ND). Relative contributions inferred from point mutations are indicated by the color coding as follows, strongest impact (<50% of wild-type binding) = coral, medium impact (50% to 90%) = light peach, minor or no effect (>90%) = light blue). **(C)** Binding results using DmORC complex (Orc1–6) expressed and purified from ExpiSf9 cells, where the complex contains either full-length wild-type Orc1 or Orc1 carrying combined multiple mutations in R1 and R2 (Mut R1/R2; see Figure 2A). **(D)** Boxes indicate the SLiMs, Region 1 and 2 (R1 and R2), identified by the deletion analysis (Figure 1C) and point mutations (Figures 2A**, 2B**) as contributing to HP1a binding; color coding is as described in (Figure 2B).

Having identified two SLiMs (R1 and R2) in Orc1 IDR required for HP1a interaction, we next sought to identify the critical residues within R1 and R2. We generated mutations contain single amino acid substitution in R1 and R2 (where alanine replaced bulky side-chain residues) and quantified their HP1a-binding relative to the WT Orc1 (2–240) fragment. The roles of individual bulky side-chain residues were revealed by analyzing the binding of Orc1 (2–240) fragments containing single alanine mutations (**Figure 2B, Supplementary Figure 3)**. Some mutated residues resulted in major reductions in HP1a binding (R200, R206, K236, and R237, pink), while others caused moderate reductions (R203, K207, and V234, yellow) and the rest displayed no significant change (P201, L209, E235, R238, and R239, blue) (**Figure 2B)**. Our results show critical residues within R1 and R2 regions of the Orc1 IDR are essential for HP1a binding.

To test if R1 and R2 were necessary for binding of HP1a by full-length Orc1 in the context of the intact ORC complex, we reconstituted the complex by overexpressing and purifying DmOrc1–6 from ExpiSf9 cell culture. Complexes formed with WT full-length Orc1 displayed robust HP1a binding, whereas those containing Mut R1 and Mut R2 bound HP1a with undetectable level (**Figure 2C**). We conclude that R1 and R2 dependency is not restricted to analyses of Orc1 fragments but are also critical for HP1a binding by ORC complex containing full-length Orc1.

We conclude that two separated SLiMs (R1 and R2) within the DmOrc1 IDR are both required for HP1a binding *in vitro*. Different residues in the elements, in particular, positive charged amino acids make critical contributions to this interaction (**Figure 2D**), but the relative spacing between R1 and R2 seems tolerate to modest alterations. The key positive charged residues in R1 and R2 (notably R200, R203; K236 and R237) are conserved across *Drosophilids* where reliable sequencing data is available (**Supplementary Figure 4**). In contrast, the length of R1 is variable among species, although most retain clusters of positively charged residues (R/K). The spacing between R1 and R2 is also variable, primarily due to differences in length and composition variations within the intervening region (residues 225–233 in Orc1), consistent with the fact our observation that some DmOrc1 spacer deletions had no strong impact on HP1a binding (**Figure 1C**).

### Orc1 binding requires HP1a dimerization, the N-terminal hexa-glutamic acid motif, and the C-terminal extension

HP1a predominantly exists as a dimer in cells, which is mediated by its C-terminal extension (CSD) ^19^ (**Figures 3A, 3B)**. The dimerization of HP1a and its C-terminal extension (CTE) region are in turn required for binding to many HP1a partner proteins ^16,20^. We thus analyzed the impact of HP1a mutants that eliminate or enhance HP1a dimerization on Orc1 interactions. An I191E mutant of HP1a, which is known to exist predominantly as a monomer ^21^, completely lost its ability to bind to the Orc1 (2–240) fragment, even when the fragment concentration was increased 2- or 4-fold (**Figure 3C**). By comparison, an HP1a construct bearing the R188Q mutation, which enhances HP1a dimerization ^22^, displayed an increased binding affinity for Orc1 (2–240) relative to wild-type HP1a (**Figure 3C**). I191 is highly conserved across many *Drosophila* species and is important for maintaining the dimerization state of HP1a, whereas R188 has frequently substituted with glutamine (Q), a change that enhances HP1a dimerization (**Supplementary Figure 5**). These observations show that in our assays HP1a dimerization is required for Orc1 binding.

**Figure 3.**
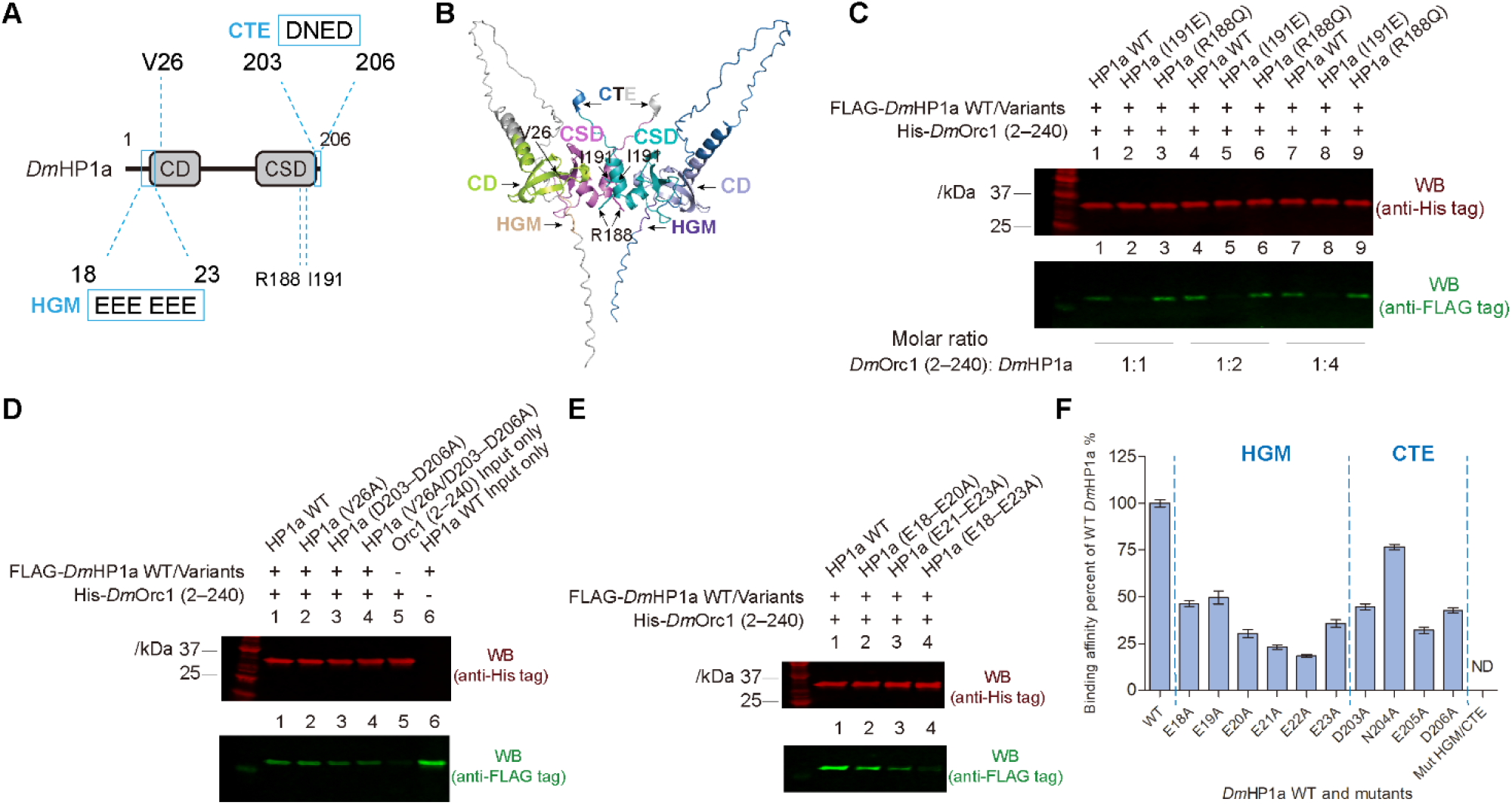
Distinct N- and C-terminal motifs (HGM and CTE) in HP1a and its dimerization mediate Orc1 binding. **(A)** Potential candidate Orc1 binding determinants within HP1a are highlighted: the N-terminal hexa-glutamic acid motif (HGM, residues E18–E23); the previously identified critical residue V26, and the C-terminal extension (CTE, DNED, residues D203–D206); Residues R188 and I191 that acts as HP1a dimerization determinants. **(B)** AlphaFold predicted structure of *Drosophila melanogaster* HP1a dimer with all potential Orc1-binding determinants labeled are shown in Figure 3A. **(C)** HP1a dimerization maintains its Orc1 binding affinity in vitro. Monomeric HP1a (I191E) and an enhanced dimer mutant (R188Q) were assayed for binding to the wild-type Orc1 (2–240) fragment across a range of molar ratios of Orc1 (2–240) to HP1a (wild-type and mutants). **(D)** Mutating the CTE (DNED) to tetra-alanine (D203–D206A) reduced Orc1 binding significantly, whereas the HP1a V26A CD mutation did not, nor did it further decrease binding in combination with D203–D206A. **(E)** Mutating all 6 glutamates in HGM to alanines (E18–E23A) reduced Orc1 binding significantly, while mutating either half of HGM (E18–E20A, E21–E23A) resulted in partial loss of Orc1 binding. **(F)** Quantification of Orc1 (2–240) binding for the indicated single alanine substitution in HGM (18–23) and CTE (203–206), displayed as a percentage of wild-type HP1a binding; all residues in HGM and CTE were mutated to alanine in Mut HGM/CTE.

Dimerization of the CSD generates a hydrophobic binding surface that recruits numerous partner protein via PxVxL motifs ^16,23^. However, the Orc1 R1 and R2 sequences are not homologous to this motif, thus we experimentally asked where within HP1 these Orc1 motifs might bind. One study suggested that the adjacent CTE (D203–D206A, **Figure 3A**) also contributes to interaction-partner specificity ^22^, and we also note that the HP1a CTE is highly conserved across *Drosophilids* (**Supplementary Figure 5).** Indeed, mutating the HP1a CTE to tetra-alanine (DNED→AAAA) caused a reduction in binding to the Orc1 (2–240) fragment compared to WT HP1a (**Figure 3D**), showing a key role for the CTE in Orc1 binding. Further binding analyses and quantification data identified D203, E205, and D206 as key residues required for the interaction, as alanine substitution mutants D203A, E205A, and D206A markedly reduced Orc1 (2–240) binding (**Figure 3F, Supplementary Figure 6**).

The dependence of Orc1 binding on HP1a dimerization suggested that the two Orc1-binding determinants, either R1 or R2 may engage another interaction motifs within an HP1a dimer rather than a single HP1a binding site. We therefore sought to identify an additional HP1a motif that cooperates with the CTE to engage Orc1 recognition. As a clue to the location of a second Orc1-binding motif, we revisited the V26 and its adjacent N-terminal region of HP1a. The natural occurring variant of *Drosophila* HP1a, V26M, was shown to disrupt gene silencing in vivo ^24^ and this substitution in the CD (V26M) has been reported to abolish its binding to synthetic peptide of H3K9me2 tail in vitro ^25^, suggesting that it may have functions beyond canonical recognition. However, neither the V26A mutation nor an expanded V26–K28A substitution (^26^VEK^28^ to ^26^AAA^28^) detectably affected binding to the Orc1 (2–240) fragment compared to the wild-type HP1a (**Supplementary Figure 7**). Furthermore, introduction of the V26A into the CTE mutant variants failed to reduce further Orc1 binding (**Figure 3D compare lanes 3 and 4**). These results imply that V26 and V26–K28 are unlikely to constitute a determinant of the Orc1–HP1a interaction.

We re-evaluated the HP1a N-terminal region surrounding V26, guided by the biochemical properties of the HP1a-binding motifs R1 and R2 in Orc1 IDR. As the critical residues for HP1a binding within both R1 and R2 are predominantly basic, we reasoned that their binding partners in HP1a might contain acidic residues. Examination of the HP1a sequence revealed a striking acidic stretch composed of six consecutive glutamate residues (18–23) located upstream of V26. An AlphaFold-predicted structural model of an Orc1–HP1a complex positions this acidic stretch in close proximity to the Orc1 IDR. Moreover, this acidic stretch (residues 18–23) spans the boundary of two previously characterized N-terminal deletions of HP1a. Deletion of residues 1–21, which removes only part of the motif, reduced ORC binding in pull down assays with *Dm* embryo extracts, whereas deletion of HP1a residues 1–31, which eliminates the motif entirely, abolished the interaction ^6^. Together, these observations identify a candidate motif within HP1 for Orc1-binding, the hexa-glutamic acid motif (**Figure 3A**, ^18^EEEEEE^23^, hereafter called ‘HGM’), which is evolutionarily conserved among *Drosophilids* (**Supplementary Figure 8**).

To test the role of the HGM, mutating either half of the HGM motif to tri-alanine (^18^AAA^20^ or ^21^AAA^23^) reduced HP1a binding to Orc1 (2–240) fragments, while no binding was detected when all 6 glutamic acids were mutated to alanines (**Figure 3E**). To define residue-specific contributions, each glutamic acid within the HGM was individually mutated to alanine, and the effects on Orc1-binding were evaluated. Residues E21–E23, located proximal to the CD, contributed more to Orc1 (2–240) binding than E18–E20 (**Figure 3F, Supplementary Figure 6**). This finding is consistent with sequence analyses showing that the acidic stretch tolerates some variation in both length and composition across species, including substitutions with neutral residues (alanine, valine, or glutamine) (**Supplementary Figure 8**). The overall composition of this region is consistent with a negatively charged cluster likely involved in electrostatic interactions.

We conclude that two distant HP1a motifs are required for Orc1 binding: the N-terminal HGM and the C-terminal CTE. Key residues in the HP1a HGM and CTE are negatively charged, suggesting that stable interactions require salt bridge formation with the corresponding positively charges residues in Orc1 R1 and R2.

### The HP1a CTE interacts with Orc1 R2, and the HGM interacts with Orc1 R1

These observations raised the question of how the Orc1 R1/R2 motifs might pair with the HP1a HGM and CTE elements. In principle, the R1 motif of Orc1 could interact with the HGM elements of HP1a and the R2 with the CTE, or *vice versa* (**Figure 4A**). We reasoned that, if the HP1a CTE specifically interacts with R2 of Orc1, then combining both mutations within CTE and R2 would not reduce binding in comparison to WT HP1a binding to the Orc1 R2 mutant. In contrast, combining CTE and R1 mutants would more severely reduce binding, since both critical regions would be disrupted. We observed that HP1a bearing a mutated CTE was further impaired in binding to a Orc1 variant lacking its R1 (Orc1 2–240, Δ201–210), whereas this HP1a mutant bound as well as wild-type HP1a to Orc1 in the absence of R2 (Orc1 2–240, Δ231–240) (**Figure 4B**). These data indicate that the HP1a CTE interacts with Orc1 R2, and the HP1a HGM interacts with Orc1 R1 (**Figure 4C**). Evolutionary conservation enhances our confidence in these quantifiable differences. The CTE and R2 are short and highly conserved across *Drosophilids*, whereas the HGM and R1 have some variance in sequence length and composition, while retaining key charged residues in both (**Supplementary Figures 4**, **5**, and **8**). Although we will introduce further notions for how this pairing may occur in the dimer of HP1 in the discussion, we favor the idea that an Orc1 protomer engages both subunits of an HP1a dimer, where one HP1 subunit provides the CTE and the other subunit provides the HGM (*trans* contacts), as opposed to a single subunit of HP1a in the dimer providing both motifs (*cis* contact).

**Figure 4.**
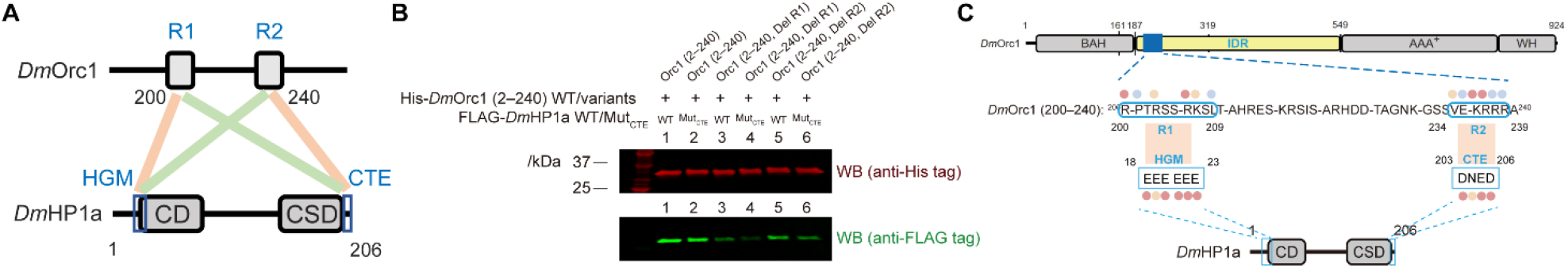
Pairing of Orc1 R1/R2 with HP1a HGM/CTE motifs. **(A)** Hypothetical pairing schemes between Orc1 and HP1a, illustrating R1–HGM and R2–CTE interactions (light peach) or R1–CTE and R2–HGM interactions (light green). **(B)** Binding results for revealing the pairing between Orc1 motifs and HP1a. The Orc1 variant Orc1 2–240, Del R1 (201–210) and Orc1 2–240, Del R2 (231–240) were used to assess the contributions of R1 and R2 interaction with CTE in HP1a. FLAG-*Dm*HP1a used in lane 4 and called Mut_CTE_ corresponds to the HP1a variant harboring the D203–D206A substitutions (DNED→AAAA; see Figure 3D). **(C)** Experimentally supported pairing model based upon the data in Figure 4B.

### Reduced Orc1–HP1a interaction suppresses heterochromatin-mediated gene silencing (PEV)

The identification of Orc1 IDR mutations that are critical for HP1a binding provided an opportunity to investigate the impact of disrupting this biochemically defined interaction upon cellular and organismal phenotypes. Two mutant alleles of *orc1* encoding proteins were introduced into the endogenous *Drosophila* locus using CRISPR-Cas9 mediating genome editing ^26^. One version deletes the essential 40 amino acid stretch (aa200–240) in IDR of Orc1 (strain called *orc1*^Del^ ^R1–R2^), the other contains point mutations within R1 and R2 of Orc1 (strain called *orc1*^Mut^ ^R1/R2^), which harbor the same mutations in DmOrc1 (2–240, Mut R1/R2) see Figure 2A (**Figure 5A**). Both mutations were viable as homozygotes, demonstrating that these residues and they mediate Orc1 binding to HP1a are not essential for organismal development, in contrast to Orc1 subunit null mutations ^27^.

**Figure 5.**
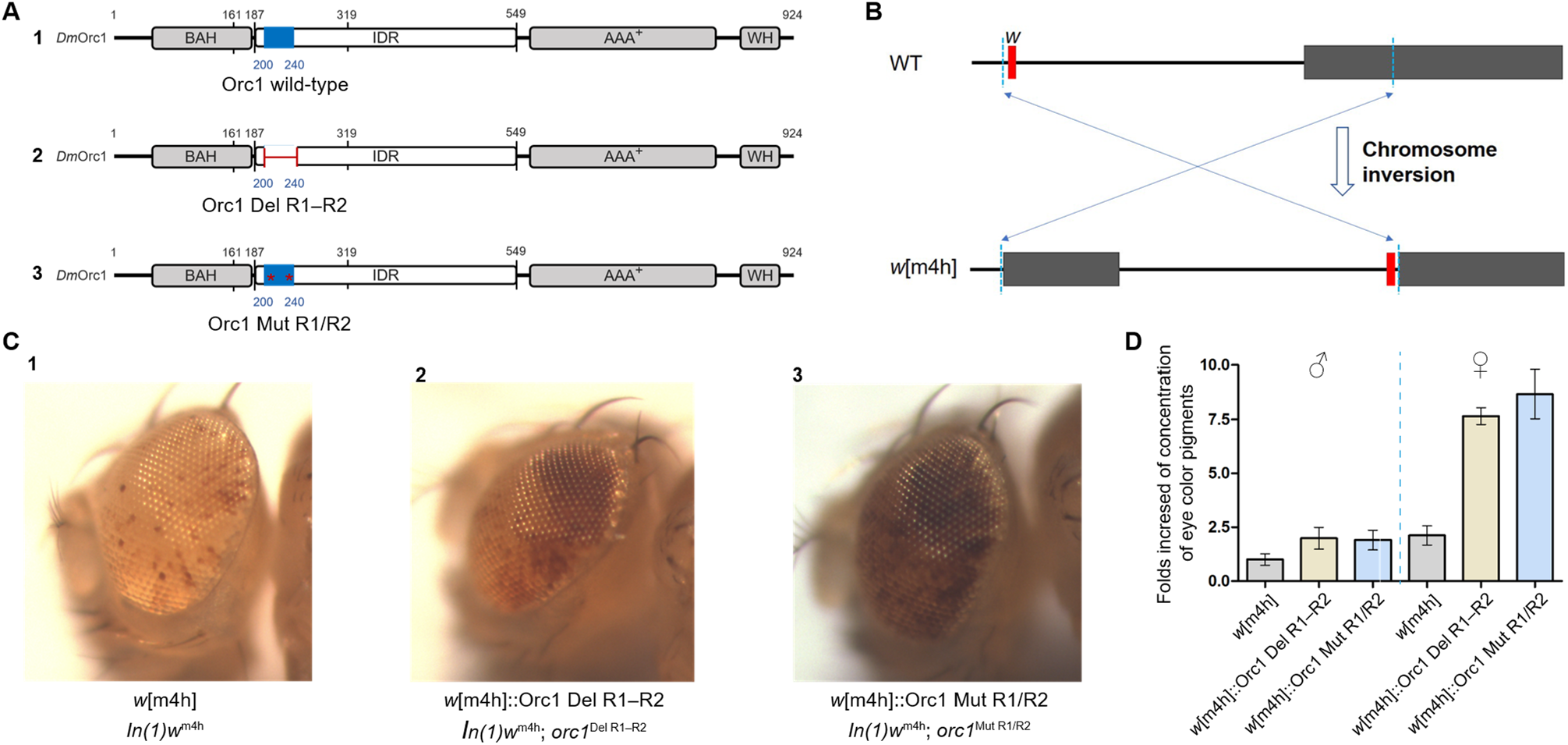
Dominant suppression of Position-Effect Variegation (PEV) in the *Dm*Orc1 mutants. **(A)** Fly strains with mutations at the endogenous *orc1* locus were generated using CRISPR-cas9 genome editing: wild-type Orc1 (**A1**), *orc1*^Del^ ^R1–R2^ deletes aa 200–240, which includes R1, R2, and the 25 amino-acid linker between (**A2**), and *orc1*^Mut^ ^R1/R2^ has all large side-chain residues in R1 and R2 mutated to match DmOrc1 (2–240) Mut R1/R2 (see Figure 2A) (**A3**). **(B)** *In(1)w*^m4h^ results in PEV of the *white*^+^ gene, caused by inversion of the *w* locus into proximity with heterochromatin. **(C)** Examples of individual eye colors of male *w*[m4h] (**C1**), *w*[m4h]::Orc1 Del R1–R2 (**C2**), and *w*[m4h]::Orc1 Mut R1/R2 (**C3**), with the indicated genotypes. **(D)** Quantification of eye pigment differences in male and female flies for the indicated genotypes. Error bars show variances from 4 different measurements. Red pigment was extracted from pooled males or females and measured by optical absorbance as described in the materials and methods.

We first focused on whether Orc1 binding to HP1a is critical for constitutive heterochromatin function by evaluating its impact on heterochromatin-mediated gene silencing (Position-Effect Variegation or PEV) of the *white* gene in *In(1)w*^m4h^ (**Figure 5B**). Both homozygotes flies harboring *orc1* mutants, *orc1*^Del^ ^R1–R2^, and *orc1*^Mut^ ^R1/R2^, respectively, exhibited dominant suppression of *In(1)w*^m4h^ PEV, resulting in eyes with extensive red patches compared to controls (**Figure 5C**). Quantification of total eye-pigment levels showed a ∼2-fold increase in Orc1 mutant males and ∼4-fold increase in mutant females, relative to controls (**Figure 5D**). We conclude that disrupting Orc1–HP1a binding results in a Su(var) phenotype, demonstrating that the interaction is required for heterochromatin-mediated gene silencing.

### Decreased Orc1–HP1a interaction leads to increased nucleolar volume and rRNA transcription

Heterochromatin-mediated gene silencing also regulates the expression of tandemly repeated ribosomal RNA genes (rDNA), whose transcription forms the nucleolus, a prominent multi-layered condensate required for ribosome assembly ^28^. In most eukaryotes, only a subset of rDNA repeats is transcribed, while the remaining copies are silenced through heterochromatin formation ^29,30^. This balance is regulated both at the chromosomal level, where rDNA repeats are embedded next to heterochromatin ^31,32^, and cell-biologically, where silent rDNA repeats are positioned outside the nucleolus ^33^ (**Figure 6A**). HP1 and H3K9me2/3 are enriched at silent rDNA in both *Drosophila* and mammals ^34,35^, and HP1 is required for maintaining nucleolar organization by excluding other repetitive DNAs from the nucleolus ^36^. Mutations in HP1 cause increased rDNA transcription, enlarged or disorganized nucleoli, and rDNA instability ^37–39^, demonstrating that nucleolar structure and function is sensitive to changes in heterochromatin.

**Figure 6.**
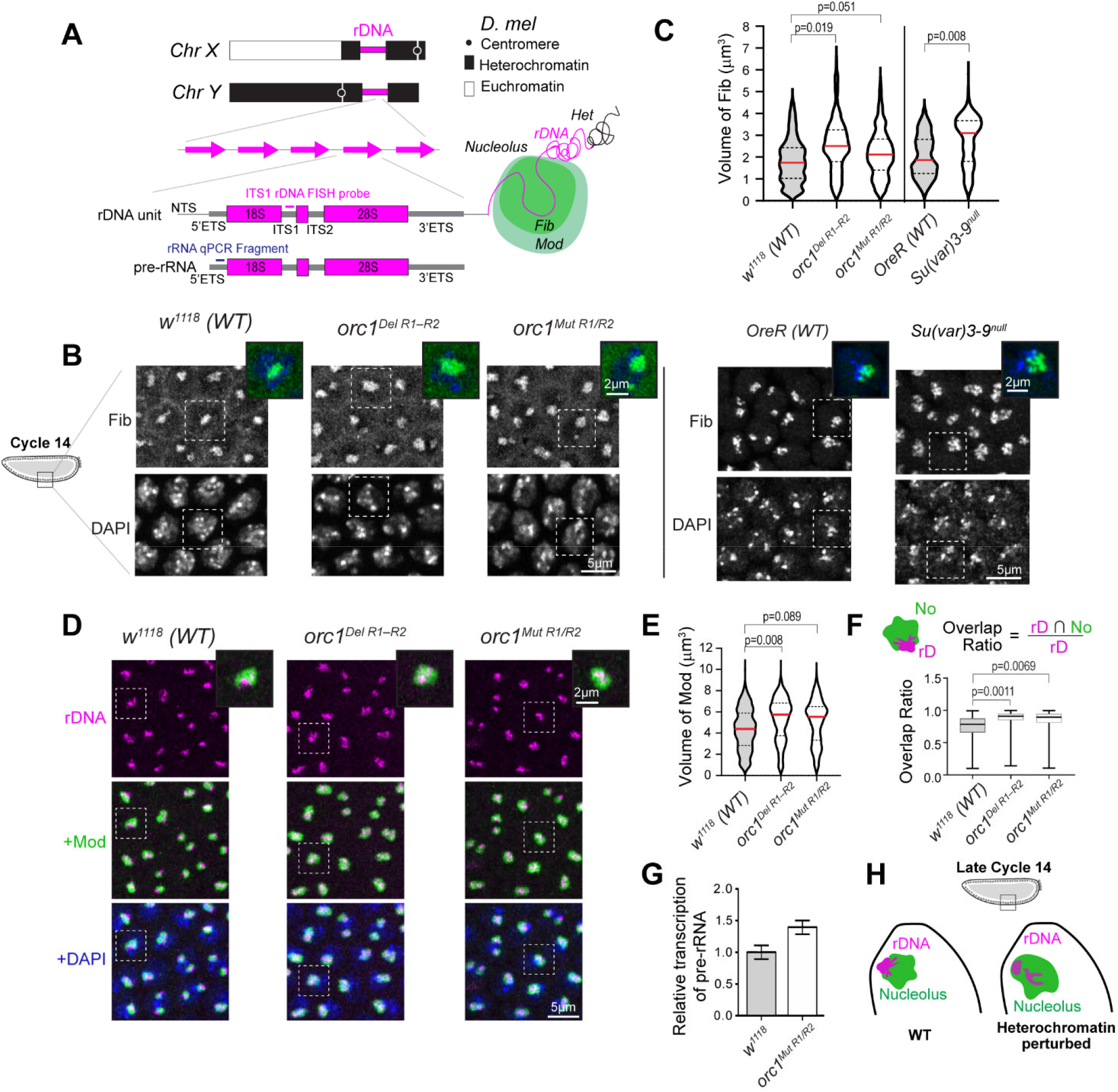
Decreased Orc1–HP1a interaction cause nucleolar expansion and rDNA derepression. **(A)** *Drosophila* ribosomal DNA (rDNA) loci are embedded within pericentromeric heterochromatin on the X and Y chromosomes. rDNA genes are arranged in head-to-tail tandem repeats, with each repeat unit consisting of a transcribed region and non-transcribed spacers (NTS). The transcribed region produces the 47S precursor rRNA (pre-rRNA), which contains external transcribed sequences (5′ ETS and 3’ ETS) and internal transcribed spacers (ITS1 and ITS2) that are removed during processing to generate mature 18S, 5.8S and 28S rRNAs. The position of the ITS1 probe used for rDNA fluorescence in situ hybridization (FISH) is indicated on the rDNA repeat unit. The amplified fragment used for pre-rRNA quantification by RT-qPCR is marked on the pre-rRNA. A subset of rDNA repeats is actively transcribed in the nucleolus, while some repeats are positioned in a high intensity block outside the nucleolus and are transcriptionally silent. The nucleolus forms a multi-layered, membrane-less compartment around actively transcribed rDNA, with the Dense Fibrillar Components (DFC) layer marked by Fibrillarin (Fib) and the outermost Granular Component (GC) visualized using Modulo (Mod) in this study. **(B)** (Left) Maximum-intensity z projections of nuclei from *w*¹¹¹⁸ control, *orc1*^Del^ ^R1–R2^ and *orc1*^Mut^ ^R1/R2^ mutant embryos at late nuclear cycle 14, stained for the nucleolar marker Fib and DNA (DAPI). *orc1* mutant embryos display enlarged nucleoli relative to controls. (Right) OreR (WT control) and *Su(var)3-9*^null^ embryos stained with Fib and DAPI at nuclear cycle 14 are shown for comparison. Scale bars, 5 µm. **(C)** Violin plots show the distribution of nucleolar volume based on Fib segmentation in late nuclear cycle 14 embryos of the indicated genotypes, with width indicating density of data points. Horizontal lines indicate median (red, solid) and quartiles (black, dashed). (Left) *w*^1118^/WT, n = 330 nucleoli from 7 embryos; *orc1^Del^ ^R1–R2^*, n = 334 nucleoli from 8 embryos; *orc1^Mut^ ^R1/R2^*, n = 358 nucleoli from 8 embryos. (Right) *OreR/wt*, n = 544 nucleoli from 7 embryos; *Su(var)3-9* ^null^, n = 455 nucleoli from 6 embryos. Statistical comparisons were performed using a linear mixed-effects model with genotype as a fixed effect and embryo as a random effect, to account for non-independence of nucleoli within the same embryo (See Methods for details). **(D)** Representative z-projections of late nuclear cycle 14 embryos of the indicated genotypes, stained for rDNA by FISH (magenta), the nucleolar marker Modulo (green), and DNA (DAPI, blue). *orc1* mutant embryos exhibit altered rDNA distribution and enlarged nucleoli compared to control. **(E)** Violin plots show the distribution of nucleolar volume based on Mod segmentation in late nuclear cycle 14 embryos of the indicated genotypes, with width indicating density of data points. Horizontal lines indicate median (red, solid) and quartiles (black, dashed). *w*^1118^/WT, n = 754 nucleoli from 15 embryos; *orc1^Del^ ^R1–R2^*, n = 669 nucleoli from 15 embryos; *orc1^Mut^ ^R1/R2^*, n = 308 nucleoli from 7 embryos. Statistical comparisons were performed using a linear mixed-effects model. **(F)** Box-and-whisker plots showing rDNA-nucleolus overlap ratio, defined as the intersection volume between segmented rDNA (rD) and nucleolar (No) signal divided by total rDNA volume (rD ∩ No / rD). Boxes represent the interquartile range with the median indicated by a horizontal line and whiskers extend from minimum to maximum values. *w*^1118^/WT, n = 376 rDNA-nucleolar pairs from 7 embryos; *orc1^Del^ ^R1–R2^*, n = 349 rDNA-nucleolar pairs from 8 embryos; *orc1^Mut^ ^R1/R2^*, n = 319 rDNA-nucleolar pairs from 7 embryos. Statistical comparisons were performed using a linear mixed-effects model. **(G)** Transcriptional levels of pre-rRNA in *Drosophila* WT and *orc1*^Mut^ ^R1/R2^ embryos at late nuclear cycle 14, measured by RT-qPCR and normalized to *rp49* mRNA levels. n=3; error bars, SD. **(H)** Schematic summary of the observed phenotypes, illustrating that reduced H3K9me2/3, HP1a levels, or Orc1–HP1a interactions lead to rDNA de-condensation, redistribution of rDNA into the nucleolus, and nucleolar enlargement.

Since Orc1 mutations decrease HP1a interactions and suppress PEV, we asked whether these mutations impact rDNA and nucleolar organization. We therefore examined nucleolar volume of *orc1* mutant embryos in late nuclear cycle 14, a stage at which heterochromatin is established following the mid-blastula transition and the embryo consists of a relatively uniform population of diploid somatic cells ^40,41^, by immunostaining for the nucleolar marker Fibrillarin (Fib). *orc1*^Del^ ^R1–R2^ and *orc1*^Mut^ ^R1/R2^ mutant embryos showed increased nucleolar volumes compared to wild-type controls (Median: *w*^1118^ = 1.747 µm^3^, *orc1*^Del^ ^R1–R2^ =2.513 µm^3^, *orc1*^Mut^ ^R1/R2^ = 2.125 µm^3^ (left panels of **Figures 6B, 6C**). To test whether this phenotype reflects disruption of heterochromatin at rDNA, we examined the impact of other prominent heterochromatin regulators on the nucleolus. Su(var)3-9 is a histone methyltransferase responsible for H3K9 methylation and heterochromatin formation ^42^. Loss of Su(var)3-9 is associated with reduced levels of H3K9me2 (**Supplementary Figures 9A, 9B**). Consistently, homozygous *Su(var)3-9*^null^ embryos derived from homozygous mutant mothers showed a 1.6-fold increase in nucleolar volume relative to wild-type controls (OreR (wt) = 1.862 µm^3^ and *Su(var)3-9* ^null^= 3.11 µm^3^) (right panels of **Figures 6B, 6C**). Similarly, maternal knockdown of HP1a by RNAi resulted in nucleolar enlargement (**Supplementary Figures 9G, 9H**). These findings demonstrate that disrupting heterochromatin components causes nucleolar volume expansion. Notably, nucleolar enlargement in *orc1* mutant embryos occurred without a detectable global loss of H3K9me2 (**Supplementary Figures 9C, 9D**), suggesting that Orc1 influences rDNA regulation independent of bulk H3K9 methylation levels, likely through direct effects on HP1a localization or stability at rDNA loci. This may also explain why *Su(var)3-9*^null^ and HP1a RNAi embryos exhibit a stronger nucleolar phenotype than *orc1* mutants, as disruption of H3K9 heterochromatin or HP1a function would be expected to have stronger effects than selectively disrupting the Orc1–HP1a interaction.

To visualize whether increased nucleolar volume reflects changes in rDNA distribution, we performed combined immunofluorescence and FISH (immuno-FISH) at late nuclear cycle 14 using a probe targeting the ITS1 region of rDNA repeats (**Figure 6A**). In wild-type embryos at this stage, the majority of rDNA signal localizes within the nucleolus, while a smaller fraction remains positioned outside, forming a distinct high-intensity block at the nucleolar periphery ^43^. This subset likely represents transcriptionally inactive, ‘heterochromatinized’ rRNA genes. Combined immuno-FISH analysis revealed that the high-intensity rDNA block is reduced in *Su(var)3-9*^null^ embryos, with corresponding increases in the amount of dispersed rDNA within the nucleolus (**Supplementary Figures 9E, 9F)**. Similar rDNA redistribution was also observed in cycle 14 embryos upon maternal knockdown of HP1a (**Supplementary Figures 9G–9I**). Both types of *orc1* mutant embryos phenocopied these outcomes, displaying reduced high intensity blocks outside the nucleolus, expanded rDNA signal inside the nucleolus, and increased nucleolar volumes (**Figures 6D–6F**). We conclude that heterochromatin components and Orc1–HP1a binding are required for normal nucleolar and rDNA organization.

The observed nucleolar phenotypes suggest that disrupting Orc1–HP1a binding impacts heterochromatin-mediated silencing of rDNA genes. To directly test whether rDNA redistribution and nucleolar enlargement are associated with increased rDNA transcription, we measured rRNA synthesis in *orc1* mutant embryos. We used RT-qPCR to quantify precursor rRNA (pre-rRNA) levels, which reflect nascent transcription rather than mature rRNA species (see methods and **Figure 6A** for strategy and amplicon locations). RT-qPCR of total RNA isolated from nuclear cycle 14 embryos revealed a 1.4-fold increase in pre-rRNA levels in *orc1*^Mut^ ^R1/R2^ mutant embryos compared to *w*¹¹¹⁸ wild-type controls (**Figure 6G**). We conclude that disrupting Orc1–HP1a interactions leads to increased rDNA transcription, redistribution of rDNA, and enlargement of nucleoli (**Figure 6H**), demonstrating another key role for Orc1–HP1a interactions in heterochromatin function.

## Discussion

The present study builds on a prior report that the Orc1 protein mediates Origin Recognition Complex (ORC) binding to Heterochromatin Protein 1a (HP1a) ^6^, by confirming that interaction and revealing the molecular components responsible for association, and their impact on heterochromatin structure and function in living organisms. First, mutational and biochemical analyses show that two SLiMs in the Orc1 IDR (R1and R2) bind two spatially distant motifs in HP1a (HGM and CTE). Orc1 R1 binds to the HP1a hexa-glutamic acid motif (HGM), while R2 binds the HP1a C-terminal extension (CTE), identifying a novel binding mechanism compared to previously described interactions between HP1a and its partner proteins ^16,23^. Second, when deletion and point mutations in R1 and R2 that disrupt Orc1–HP1a interactions were generated at the endogenous *orc1* locus cause loss of heterochromatin-mediated gene silencing, rDNA de-repression, and nucleolar enlargement without loss of viability. Deletion of the entire Orc1 is not viable (Parker et al 2019) and other such SLiMs within this IDR, as for the HsORC1 IDR ^44^ may be required for essential DNA replication functions. We conclude that Orc1–HP1a interaction is required for heterochromatin functions.

### A novel mechanism for Orc1**–**HP1a binding

The two HP1a-binding motifs, R1 ^200^RPTRSSRKSL^209^ and R2 ^234^VEKRRR^239^, respectively, locate at the opposite ends of a 40-amino acid segment within the Orc1 IDR (**Figure 2D**). Similarly, the two corresponding motifs in HP1a required for this interaction are located in distant regions of the protein. The HGM (residues 18–23, EEEEEE), which interacts with Orc1 R1, is located within the N-terminal extension (NTE) of HP1a, whereas the CTE (residues 203-206, DNED), which interacts with R2 (**Figure 4C)**, is positioned 180 amino acids away. While decreasing the 25 aa spacing between R1 and R2 did not significantly affect HP1a binding (**Figure 1C**), single amino acid mutations within R1 and R2, especially the positive residues and in combination did abolish Orc1–HP1a interactions (**Figure 2B**). We propose that the critical positive residues in Orc1 R1 and R2 bind to the critical negative residues in HP1a HGM and CTE through salt bridges. Arginine residues in Orc1 R1 pairing with the hexa-glutamic acid sequences in HP1a, while the ^236^KRRR^239^ amino acid stretch in Orc1 R2 pairing with the ^203^DNED^206^ residues in HP1a (**Figure 4C**). These critical R/K residues within R1 and R2 of Orc1 IDR, HGM-like acidic stretch within NTE of HP1a, the CTE of HP1a, and HP1a dimerization determine residues (**Supplementary Figures 4, 5**) are conserved across 40 MY of *Drosophila* evolution, implying that the Orc1 interactions with HP1a is subject to selection.

The observed mode for HP1 binding by Orc1 is distinct from previously known examples. Most of the 100s of HP1 binding partners identified across many species, from fission yeast to humans, utilize a PxVxL binding motif, suggesting its functional and evolutionary importance ^20,23,45,46^. Structural studies have shown that PxVxL peptides bind asymmetrically to a hydrophobic pocket formed by the HP1 CSD dimer, with the central Val occupying the CSD dimer C-terminal cleft ^20^. A recent study comprehensively defined *Drosophila* HP1 Access Codes (or HACs), and demonstrated the insufficiency of PxVxL motifs for HP1 binding. Specifically, residence in IDRs, positive residues flanking degenerate PxVxL motifs, and interactions with the HP1 CTE are required for effective binding ^16^. Although the Orc1 R2 motif also binds to the HP1 CTE, our finding that Orc1 lacks PxVxL motifs and binds to the distant HMG of HP1a requiring a distinct positively charged R1 motif demonstrates that this binding mode is distinct from the mechanism used by HACs. Additional studies are needed to determine if the Orc1 binding mechanism described here is unique, or shared with other HP1 partners that do not present the canonical PxVxL sequence.

### Why do Orc1**–**HP1a interactions require HP1 dimerization?

HP1 predominantly forms homodimers in vivo and in vitro, mediated by interactions between each monomer’s chromoshadow domain (CSD) ^19^. We showed that the HP1a dimer is the preferred binding partner for Orc1, since CSD mutations that disrupt HP1a dimerization abolish Orc1–HP1a binding, while those that stabilize the dimer enhance the Orc1 interaction (**Figure 3C**). The critical role of dimerization has important implications for understanding the geometry of Orc1 – HP1 complex and the binding stoichiometry. Homodimerization could be required because the physical separation of the CTE and the HGM motifs in an HP1 monomer is incompatible with binding to both Orc1 R1 and R2. For example, in the simplest case, one Orc1 molecule could bind one HP1 dimer, in two possible configurations. If binding in *cis* were to occur, R1 and R2 from one Orc1 molecule would contact the HGM and CTE present in the same HP1a molecule. By contrast, if in *trans* binding configuration, R1 would contact the HGM of one HP1a while R2 would contact the CTE of the other HP1a (**Figure 7A, B**). Based on a cryo-EM derived structure of a human HP1α dimer bound to a nucleosome ^47^, the distance between residue V22 in ordered protein region closest to the disordered HGM and the residue E179 in CTE are greater than 50–55 Å apart in the *cis*-binding configuration, but only separated by ∼30 Å in a *trans*-binding mode (**Supplementary Figures 10A, 10B**). The furthest polypeptide chain distance that could bridge R1 to R2 is ∼84 Å (the 24 amino-acid spacer region × 3.5 Å/residue in a fully extended conformation); however, this spacer readily tolerates shortening to a maximal distance of 50 Å (through 10 residue internal deletions, **Figure 1C**) with little impact on binding. It’s hard to estimate the distance between R206 (R1) and K236 (R2) due to the dynamic nature of IDR and lacking of experimental structural data of Orc1–HP1a complex; using one AlphaFold-predicted structure of DmOrc1 (1–319), one possible distance between R206 (R1) and K236 (R2) would be ∼ 40 Å (**Supplementary Figure 10C**). Based on these spatial constraints and the requirement for a dimer HP1a, we favor a *trans*-binding model (**Figure 7A**).

**Figure 7.**
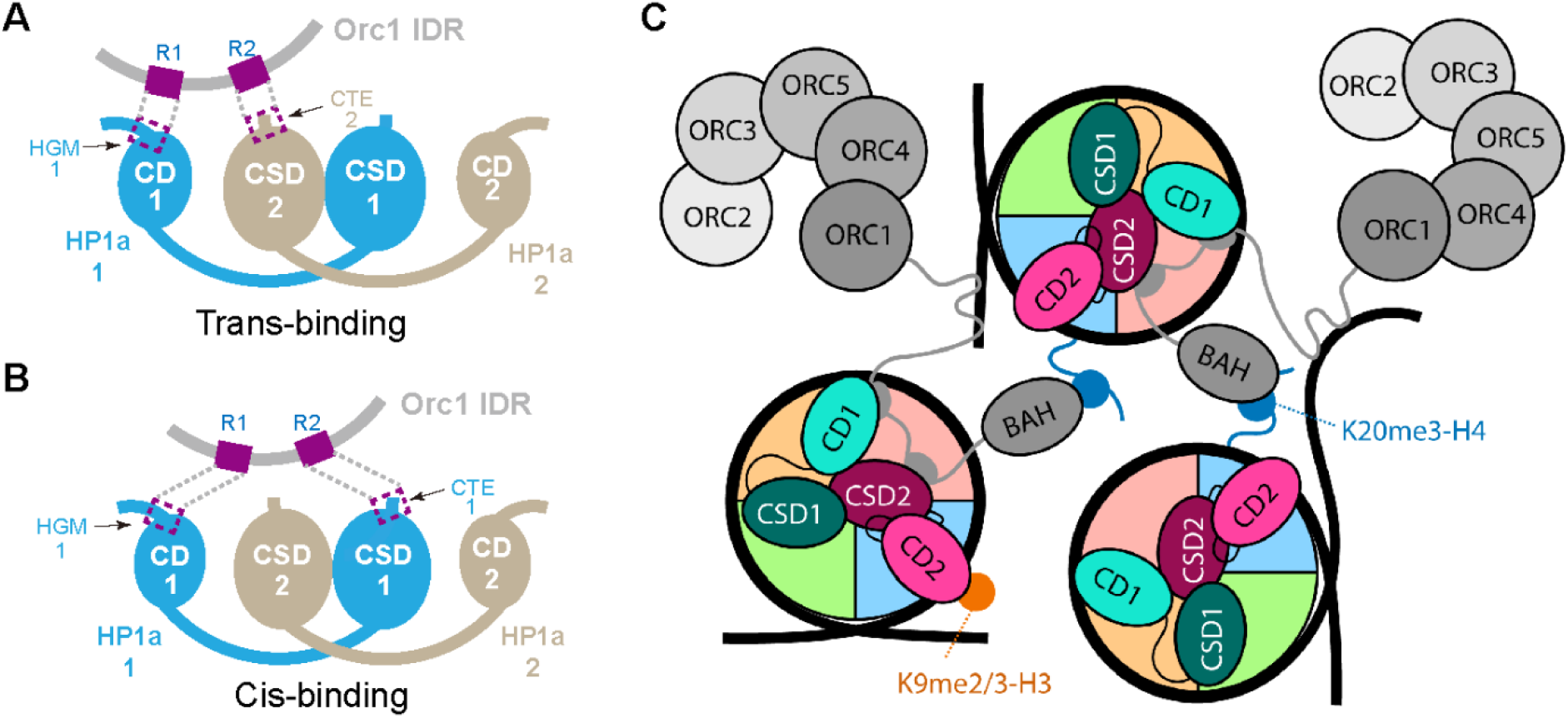
Proposed model for multivalent ORC–HP1, HP1–HP1, and ORC/HP1–histone/nucleosome interactions may facilitate nucleosome bridging and clustering, and heterochromatin organization. **(A)** This drawing models a trans binding between Orc1 and HP1a dimer, in which R1 and R2 engage motifs from different subunits of an HP1a dimer. **(B)** The model for a cis binding between Orc1 and HP1a dimer. In this model R1 and R2 engage the HGM and CTE motifs within the same HP1a subunit of a dimer. The dotted line indicates potential interaction that may be facilitated by conformational flexibility of the hinge region between the chromodomain (CD) and chromoshadow domain (CSD). **(C)** We illustrate a robust model for how multivalent ORC–HP1a interactions could mediate heterochromatin organization. ORC associates with nucleosomes through multiple interactions, including binding of the Orc1 IDR (dark gray) to motifs close to the chromodomain (CD) and chromoshadow domain (CSD) of HP1 (light/dark teal and magenta, this work), as well as to DNA (thick black lines) ^60^. In addition, the Orc1 BAH domain bind di- and tri-methylated H4K20 (dark blue) ^61^. HP1a simultaneously binding to H3K9-methylated nucleosome through its chromodomain (CD) and to DNA through its hinge region. Together, these multivalent interactions bridge neighboring nucleosomes, thereby promoting heterochromatin condensation and facilitating the spreading of heterochromatin. Nucleosome-associated HP1α dimers are schematically depicted in a conformation observed by cryo-EM (PDB ID: 8UXQ) ^47^. The cryo-EM model used is with a human HP1α homologue bound to an H2AZ nucleosome where the contacts that HP1 makes with the nucleosome do not occupy resides required for the Orc1 IDR interactions with HP1 and hence not interfering with ORC binding.

In vivo, the HP1 CSD and dimerization are required for multiple aspects of HP1’s structures and functions, including heterochromatin formation ^20,48,49^, HAC partner recruitment, HP1 oligomerization, conformational changes, interactions with histone H3 ^50,51^, and CD function ^52^. Further, heterochromatin domains behave as membraneless biocondensates formed by phase separation or similar mechanisms ^14,15^, with a key role for HP1 multivalency ^12,13^. Importantly, HP1 dimerization is absolutely required for condensate formation in vivo and in vitro ^53^. ORC also forms membraneless biocondensates in vitro, which are dependent on the same Orc1 IDR ^9^ required for the biochemical association of ORC and HP1. These observations raise the possibility that one or more of these structural and functional impacts of dimerization are required for Orc1 binding in the cell. For example, binding may not be restricted to one HP1 dimer and one Orc1 monomer, and instead rely on multivalent, dynamic interactions among multiple Orc1 proteins and HP1 dimers that coexist in solutions and condensates (**Figure 7C**). In this model, the cis and trans configurations and accompanying spatial constraints would not be dominating, since Orc1 R1 and R2 motifs could interact with HGMs and CTEs from different HP1 dimers (**Figure 7C**). This heuristic model shows a unit cell where dwell times for many of the contacts are likely short and may flicker above and below the unit to allow for compaction. Resolving the exact mode of interaction between Orc1 R1/R2 and the HGM/CTE motifs in HP1a dimers in a nucleosomal and condensate contexts will require more comprehensive structure/function analyses both within in vitro and in vivo.

### Orc1 binding to HP1a is required for heterochromatin functions

In *Drosophila*, the epigenetic state of constitutive heterochromatin is established during early embryogenesis. Heterochromatin is largely absent during the rapid cleavage divisions and emerges as replication slows, in a process dependent on the H3K9 methyltransferase Eggless/SetDB1 ^54,55^. Starting in nuclear cycle 11 and continuing through the major wave of zygotic genome activation in cycle 14, HP1a becomes enriched at H3K9-methylated nucleosomes, promoting heterochromatin condensate formation ^12,56^. Thus, these early embryonic stages represent a developmental window when heterochromatin-mediated regulatory mechanisms are established.

To dissect the *in vivo* consequences of disrupting Orc1–HP1a binding, we generated *Drosophila* strains carrying endogenous *orc1* mutations or deletions in R1 and R2. While *orc1* and *HP1a* null mutants are inviable as homozygotes ^27,57,58^, both HP1a binding-deficient *orc1* lines (*orc1*^Del^ ^R1-R2^ and *orc1*^Mut^ ^R1/R2^) are viable and fertile as homozygotes, with no detectable defects in embryonic timing or later development. Loss of Orc1–HP1a binding therefore does not appear to disrupt essential ORC-mediated replication functions, thus any observed phenotypes are not due to dysfunction in replication.

We demonstrate that Orc1–HP1a binding is, however, required for two important heterochromatin functions. First, both *orc1*^Del^ ^R1-R2^ and *orc1*^Mut^ ^R1/R2^ flies strongly suppress the Position-effect variegation (PEV) associated with a well-characterized X chromosome inversion that juxtaposes the *white* gene with pericentromeric heterochromatin (**Figure 5B**). PEV suppression is a classical phenotype observed in flies containing mutations in known heterochromatin-associated proteins, including HP1 and Su(var)3–9 ^42^. Second, Orc1–HP1a binding is required for normal heterochromatin-mediated regulation of rDNA and nucleolar organization. Heterochromatin surrounds nucleoli in diverse eukaryotes, which in *Drosophila* is mediated by the dead-box RNA helicase Pitchoune, a dual-affinity protein that is localized to the outer nucleolus and interacts weakly with HP1a through a HAC ^43^. Eukaryotic genomes, including *D. melanogaster*, contain 100s of rDNA copies that are transcribed to form the nucleolus. However, not all rDNA repeats are expressed, as a subset is silenced by heterochromatin (Guetg et al., 2010). We show that disruption of heterochromatin regulators, including Su(var)3-9 mutation or HP1a depletion, causes nucleolar enlargement and shifts rDNA localization toward the nucleolar interior during nuclear cycle 14, a developmental window when heterochromatin is being established (**Figure 6; Supplementary Figure 9**). Importantly, Orc1 mutants that disrupt HP1a-binding also exhibit nucleoli enlargement, decondensation/repositioning of rDNA to internal nucleolar locations, and rRNA transcription increases (**Figure 6**), consistent with loss of heterochromatin-mediated rDNA silencing in *Drosophila* embryos.

Overall, we conclude that Orc1–HP1a binding is required for heterochromatin functions, specifically PEV, rDNA organization and transcriptional silencing, and regulation of nucleolar size. Importantly, Orc1 R1/R2 mutations do not appear to block the establishment of heterochromatin globally, since bulk H3K9me2 levels remain largely unchanged by immunofluorescence (**Supplemental Figures 9C, 9D**). The lower magnitude of the nucleolar phenotypes in Orc1 R1/R2 mutants (*orc1*^Del^ ^R1-R2^ and *orc1*^Mut^ ^R1/R2^) compared to complete loss of heterochromatin in HP1a or Su(var)3-9 mutants, along with the retention of normal H3K9me2 levels in Orc1 R1/R2 mutants embryos, indicates that Orc1–HP1a binding is not required to establish H3K9me-marked heterochromatin. Instead, we propose that Orc1–HP1a interactions contribute to robust silencing, higher-order chromatin organization, or increasing HP1a multivalent interactions with chromatin. We speculate that Orc1, and perhaps co-recruitment of other members of the ORC complex, could promote HP1a functions in heterochromatin by aiding chromatin compaction, a property classically-associated with heterochromatin domains and gene silencing ^11,52^. For example, as noted in **Figure 7C**, HP1 dimers contain two CDs that could ‘crosslink’ H3K9me2/3 nucleosomes, perhaps facilitated by hinge-mediated interactions with DNA and unmodified nucleosomes ^59^. ORC binding to HP1 dimers could increase or strengthen chromatin compaction through the ‘reader’ activity of Orc1’s BAH domain, which binds to another heterochromatin-enriched epigenetic modification (H4K20me2/3). Alternatively, or in parallel, Orc1 binding to chromatin and HP1 could impact the formation, dynamics, or functions of HP1a-mediated heterochromatin condensates. Both Orc1 and HP1a proteins can phase separate *in vitro* and HP1a display liquid-like material properties *in vivo* ^9,12,13,53^. In this context it is particularly interesting that ORC phase separation requires the Orc1 IDR in vitro ^9^. We speculate that Orc1 or ORC binding may alter HP1 condensate composition, impact its biophysical properties such as liquidity, alter the permeability of competing factors, and/or enhance recruitment of essential heterochromatin components. The potential mechanisms for ORC-mediated regulation of heterochromatin functions are not mutually exclusive with the compaction model show in **Figure 7C**, and future experiments should focus on distinguishing between these and other models.

## Acknowledgements

We thank the work by Maren Bell who guided our designs for the CRISPR mutations and obtaining appropriate fly strains for the projects and Don Rio for discussions on our models. This work was supported by the National Institutes of Health (5R01GM141045-04) to Michael R. Botchan, the Karpen lab acknowledges support from the National Institutes of Health (R35GM139653), and James Berger the National Institutes of Health grant R35CA263778.

## Author contributions

All authors contributed to the design, writing and execution of the experiments.

## Materials and Methods

### Methods details

#### Plasmid construction and mutagenesis

Recombinant *Drosophila* N-terminal His-tagged Orc1 fragment and its truncation variants were expressed and purified from *Escherichia coli* BL21(DE3) by cloning into modified QB3 Macrolab vector 1-B contains an N-terminal 10-His tag. The coding sequence for FLAG-tagged full-length *Drosophila* HP1a was cloned into the QB3 Macrolab vector 1-B which express an N-terminal hexa-His tag and tobacco etch virus (TEV) protease-cleavage site, and overexpressed and purified from *E. coli* BL21(DE3).

Plasmids with mutant *Dm*Orc1/*Dm*HP1a were generated following site directed mutagenesis using PrimeSTAR GXL DNA Polymerase (Takara). After PCR amplification, the amplified DNA products are treated with *Dpn*I enzyme at 37°C for 30 min. After *Dpn*I treatment, the PCR products were purified QIAGEN PCR purification kit and transformed into high efficiency 5-alpha competent *E. coli* (NEB). Colonies were screened for positive mutants by plasmid DNA sequencing. Oligonucleotide sequences used to generate all of plasmids are available upon request.

### Expression and purification of recombinant proteins

#### Protein overexpression and purification of His-tagged *Dm*Orc1 fragments and FLAG-tagged *Dm*HP1a from *Escherichia coli*

N-terminal His-tagged *Dm*Orc1 and FLAG-tagged *Dm*HP1a were overexpressed using *E. coli* BL21 (DE3) as host. The His-tagged *Dm*Orc1 fragments and related mutated variants were expressed and purified using HisTrap™ HP His tag protein purification columns (Cytiva, # 17524801). Briefly, sequencing right plasmids were purified and transformed in BL21(DE3) competent *E. coli* (NEB, C2527H), transformed bacterial colonies were picked up and cultured in LB medium at 37°C overnight, and then inoculate into LB medium and culture at 37°C until the OD_600nm_ of the cell culture reaches 0.4–0.8 and then were induced to express protein with 0.5 mM of IPTG at 16 °C for 16 hours. the cultured bacteria were centrifuged to form pellet, washed, and lysed with sonication in the lysis buffer containing (20 mM Tris-HCl pH 7.5, 500 mM NaCl, 10% Glycerol), 1 cOmplete Mini Protease cocktail inhibitor tablets (Roche, #11836153001). The lysed bacterial cells were centrifuged and the supernatant is applied to the HisTrap™ HP His tag protein purification columns and gradient eluted with washing buffer contain 500 mM imidazole solution. Next, the sample was purified again with HiTrap^TM^ HP Heparin column (Cytiva, #17040701), and Gel filtration using HiPrep^TM^ 16/60 Sephacryl^TM^ S-300 HR column. All *Dm*Orc1 mutated constructs were purified according to the same experimental procedure described here. Final purified His-tagged *Dm*Orc1 were preserved within stock solution (50 mM HEPES pH 7.5, 300 mM Potassium L-glutamate, 10% Glycerol).

N-terminal His-tagged-TEV-FLAG-*Dm*HP1a and its mutants were purified with HisTrap^TM^ HP His tag protein purification columns first and then with Hi Trap^TM^ HP Heparin column. The purified His-tagged-TEV-FLAG-*Dm*HP1a were subject to purified His-tagged TEV protease to remove the N-terminal His-tag, resulting purified N-terminal FLAG-*Dm*HP1a. Then purified again with HisTrap^TM^ HP His tag protein purification column, HiTrap^TM^ HP Heparin column, and Gel filtration, respectively. Final purified N-terminal FLAG-tagged *Dm*HP1a were preserved within stock solution (50 mM HEPES pH 7.5, 300 mM Potassium L-glutamate, 10% Glycerol). Expression and purification of N-terminal His-tagged TEV protease were conducted according to the protocol ^62^

#### Expression and purification of *Dm*ORC WT and *Dm*ORC (Mut R1/R2)

Expression and purification of the *Drosophila melanogaster* ORC1–6 complex (*Dm*ORC) and the corresponding complex contains the Orc1 with mutated R1 and R2 (*Dm*ORC Mut R1/R2) were performed as previously described experimental procedures ^9^.

Recombinant baculoviruses are generated by transfecting ExpiSf9 cells with DH10Bac-derived bacmid DNA using ExpiFectamine™ Sf Transfection Reagent (Thermo Fisher Scientific). Wild-type and mutant *Dm*ORC complexes were expressed in ExpiSf9 cells using the baculovirus expression system.

For reconstitution of full-length wild-type *Dm*ORC, two separate recombinant baculoviruses were used for co-infection: one encoding *Dm*Orc1–5, in which *Dm*Orc1 carried N-terminal His6 tag and *Dm*Orc4 carried N-terminal TEV-cleavable His6 and MBP tags, respectively, and a second encoding *Dm*Orc6.

To generate the full-length mutant *Dm*ORC complex, point mutations were introduced into the R1/R2 regions of *Dm*Orc1 within the *Dm*Orc1–5 bacmid construct. The resulting baculovirus, encoding mutant *Dm*Orc1 with R1/R2 mutation together with *Dm*Orc2–5, was co-infected with a second baculovirus encoding *Dm*Orc6 to produce the full-length *Dm*ORC Mut R1/R2 complex in High5 cells.

#### Binding assay between His-tagged *Dm*Orc1 and FLAG-tagged *Dm*HP1a

To assess direct binding between *Dm*Orc1 and *Dm*HP1a *in vitro*, purified N-terminal His-tagged *Dm*Orc1 fragments and N-terminal *Dm*HP1a were used. The pull-down assays we have used with tagged and magnetic beads do not provide equilibrium measurements and we anticipated that required residues for binding within the Orc1 IDR with HP1 would be weak given the entropic penalty likely lost through specific binding. This was borne out as attempts to use the mass photometry method ^17^ were not successful.

The recombinant purified His-tagged *Dm*Orc1 fragments (10 μM) were mixed and incubated with His magnetic beads in binding buffer (20 mM HEPES pH 7.5, 100 mM KCl, 1 mM DTT, 0.5 mM PMSF, 0.05% NP-40). Then purified FLAG-tagged *Dm*HP1a (10 μM) with same binding buffer was added and mix with beads absorbed His-*Dm*Orc1 fragments, and incubated at 4°C with gentle rotation overnight to allow complex formation. The protein complex bound to beads were washed with binding buffer and the eluted with elution buffer (20 mM HEPES pH 7.5, 100 mM NaCl, 1 mM DTT, 0.5 mM PMSF, 0.05% NP-40, 500 mM imidazole). Eluted samples were resolved to 4-20% Mini-PROTEAN TGX gradient gels (Bio-Rad) and then transferred to 0.2 µm PVDF membrane using Trans-Blot Turbo Transfer System (Bio-Rad). The volume of sample applied in WB experiments with different purpose were adjusted according to different purposes.

The FLAG-*Dm*HP1a was detected using mouse anti-FLAG antibody and visualized at 800 nm using Goat anti-mouse IRdye 800CW while His-*Dm*Orc1 fragments were detected using rabbit anti-His-tag antibody and visualized at 700 nm using Goat anti-Rabbit IRdye 680RD. Imaging was performed using a Li-COR Odyssey system. Quantification was performed using Li-COR Image Studio software. All antibodies were prepared in a solution of Milk dissolved in TBST and diluted according to manufacturer’s instructions.

#### Genome editing of *Drosophila melanogaster* via CRISPR-Cas9 method

Point mutations within R1 and R2 regions of *Dm*Orc1, as well as a deletion encompassing both R1, R2 regions, and the linker region between them of *Dm*Orc1, were introduced at the endogenous *orc1* locus of *Drosophila melanogaster* using CRISPR-Cas9-mediated genome editing followed by homology-directed repair (HDR) with donor templates containing the desired modifications, as previously described ^26^. This method combines two plasmids, the first one is pCFD5 (DGRC, #1529) which is used to expressing the guide RNAs, the second is pHD-ScarlessDsRed (DGRC, #1364) that is used as donor of template. We first scanned the IDR region of interest to find and then eliminate potential off target homologies where the chosen guide RNA might create breaks and unwanted deletions by creating neutral codon changes-not changing amino acids-in the WT and Orc1 Mut R1/R2. To conduct the point mutation within both R1 and R2, and delete the region contains R1 and R2, two guide RNAs were designed. gRNA-01-R1 of *Dm*Orc1 (5’-GCGAGATGAGCGGGTTGGCC-3’), recognizing location within R1 of *Dm*Orc1 and that is complementary to the underlined part of coding region from R200 to R206 of *Dm*Orc1 (5’-A<u>GGCCAACCCGCTCATCTCGC-3’</u>). gRNA-02-R2 (5’-ACGTCGTCGTTTTTCCACTG-3’), recognizing location within R2 of *Dm*Orc1 and that is complementary to the underlined part within the coding region from S233 to R239 of *Dm*Orc1 (5’-T<u>CAGTGGAAAAACGACGACGT</u>-3’). gRNA-01-R1 and gRNA-02-R2 were cloned (introduced) into pCFD5 (DGRC, #1529) and get pCFD5-gRNA1-mR1::gRNA-mR2 of *Dm*Orc1.

The two homology arms were amplified by PCR from the genomic DNA of *nos*-*Cas9* stock (BDSC, #54591). The 5’ homology arm containing *orc1* genomic region (776 base pair upstream of *orc1* gene and 925 bp at the 5’ end of *orc1* gene region that contains region encoding the R1 and R2, 1701 bp in total), and the 3’ homology arm (1455 bp of 3’ end of the *orc1* gene region and its 47 bp downstream region, 1502 bp in total) were cloned into pHD-ScarlessDsRed (DGRC, #1364). Then the resulting vector was used as template to construct the template vector for HDR. The regions for encoding for the Orc1 mutated R1/R2 and Orc1 deleted R1–R2, respectively, were introduced into the repair template for HDR that contains 1701 bp of 5’ homology arm and 1502 bp of 3’ homology arm cloned from wild type genomic DNA. The mixture of vectors for gRNA expression and repair plasmid was injected into embryos of the *nos-Cas9* stock and screened for positive transformation using the DsRed marker by Genetivision. Positive lines with DsRed were genetically crossed to remove the *nos-Cas9* chromosome. The DsRed marker is then excised from genome by PiggyBac transposase *via* crossing to the fly stock (BDGC, #8283) that contains the PiggyBac transposase. Mutant stocks were constructed and validated by PCR and Sanger sequencing the *orc1* gene from the genomic DNA from both Orc1 Mut R1/R2 and Del R1–R2 strains. the *orc1* alleles confirmed the precision of the replacements. The two different strains with altered *orc1* alleles had in one case a deletion of the codons for the entire 40 amino acid stretch in part of the long 362 amino acid IDR encoding both the HP1 binding motifs R1 and R2 mutations (Orc1 Del R1–R2). While the other strain had the point mutations in both R1 and R2 (Orc1 Mut R1/R2). Both (Mut R1/R2 and Del R1–R2) were generated using the same method. Primers to generate homology arms and mutate and delete R1/R2 are available upon request.

#### *Drosophila* strains, genetics, and Position-effect variegation analysis

All *Drosophila* stocks were maintained under standard laboratory conditions and raised on standard Bloomington medium.

The *In(1)w*^m4h^ stock was kindly provided by Dr. Keith A. Maggert from the University of Arizona. The balanced *In(1)w*^m4h^ stock (*In(1)w^m4h^*; *Sco*/*CyO*; +/+) was generated with the assistance of Jiahe Wang in lab through multiple rounds of crossing *In(1)w*^m4h^ flies to *w*^1118^; *Sco*/*CyO*; +/+ flies.

*Drosophila* that contains both *w*^m4h^ and *orc1* with mutations (*orc1*^Mut^ ^R1/R2^ or *orc1*^Del^ ^R1–R2^) were generated by crossing described as follows. Cross *w*^m4h^; *Sco*/*CyO*; +/+ female to *w*^1118^/Y; *orc1*^Mut R1/R2^ or *orc1*^Del^ ^R1–R2^; +/+ male to get offsprings and select the *w*^m4h^/Y; *orc1*^Mut^ ^R1/R2^ or *orc1*^Del^ ^R1–R2^/*CyO*; +/+ male. Individual Curly *w*^m4h^/Y; *orc1*^Mut^ ^R1/R2^ or *orc1*^Del^ ^R1–R2^/ *CyO*; +/+ male was backcrossed to *w*^m4h^; *Sco*/*CyO*; +/+ female to get offsprings and select the virgin of *w*^m4h^; *orc1*^Mut^ ^R1/R2^ or *orc1*^Del^ ^R1–R2^/*CyO*; +/+ female. Stocks that contain *w*^m4h^ and homozygous *orc1*^Mut^ ^R1/R2^ or *orc1*^Del^ ^R1–R2^ (*w*^m4h^; *orc1*^Mut^ ^R1/R2^ or *orc1*^Del^ ^R1–R2^; +/+) were generated by crossing virgin of *w*^m4h^; *orc1*^Mut^ ^R1/R2^ or *orc1*^Del^ ^R1–R2^/*CyO*; +/+ female to *w*^m4h^/Y; *orc1*^Mut^ ^R1/R2^ or *orc1*^Del^ ^R1–R2^/*CyO*; +/+ male. The effect of reduced association between Orc1 and HP1a on *w*^m4h^ PEV were characterized the red patches from eyes of *w*^m4h^/Y male, *w*^m4h^/Y; *orc1*^Mut^ ^R1/R2^; +/+ male, and *w*^m4h^/Y; *orc1*^Del^ ^R1–R2^; +/+ male recorded by photographing, and quantified by measuring the red eye pigments at 480 nm from *w*^m4h^/Y male, *w*^m4h^/Y; *orc1*^Mut^ ^R1/R2^; +/+ male, and *w*^m4h^/Y; *orc1*^Del^ ^R1–R2^; +/+ male and *w*^m4h^ female, *w*^m4h^; *orc1*^Mut^ ^R1/R2^; +/+ female, and *w*^m4h^; *orc1*^Del^ ^R1–R2^; +/+ female, respectively. The content of eye red pigments from adult eyes were estimated according to the previously described methods ^63,64^ with minor modifications as following. Red pigments were extracted from pools of four freshly dissected heads by homogenization in 250 μl of 30% (v/v) ethanol (acidified to pH ∼2 with HCl), followed by tumbling for 20 h at room temperature. The extracts were cleared by centrifugation for 1 min at 13 000 × *g*, and their absorbance at 480 nm measured against that of extracts prepared in parallel from *w*^1118^ fly heads, which were used as blank controls.

To analyze nucleolar phenotypes in *Su(var)3-9*^null^ embryos, we crossed *Su(var)3-9*^06^/*TM3,Ser* virgins to *Su(var)3-9*^17^ /*TM3,Ser* males. After selecting *Su(var)3-9*^06/17^ adults in the F1 generation, we intercrossed these siblings to collect *Su(var)3-9*^06/17^ maternal null embryos. To knockdown HP1a in *Drosophila* embryos, we crossed matα-GAL4 (BDSC, #7063) females to HP1a RNAi males (BDSC, #33400) to generate an F1 generation with HP1a depleted in the female germlines. F1 siblings were crossed to collect embryos depleted for HP1a, most of which exhibit severe developmental defects (Park et al. 2019). We selected embryos that reached nuclear cycle 14 for analyses of nucleolar morphology.

#### Reverse Transcriptase Real Time PCR (RT-qPCR)

Total RNAs were extracted from *Drosophila melanogaster* embryos at nuclear cycle 14 using TRIzol reagent (Invitrogen, # 15596) and further purified with the Monarch Total RNA Miniprep Kit (NEB, #T2010) according to the manufacturer’s manual. Reverse Transcriptase reaction was used to synthesize the cDNA contains pre-rRNA, and *RpL*32 (*rp49*) which was used as housekeeping gene for RT-qPCR. Gene-specific reverse primers were used to generate cDNA corresponding to pre-rRNA (ETS-18S region) and the reference gene *RpL32* (*rp49*). The primers, ETS-18S-cDNA-R (5’-ATACGCTATTGGAGCTGGAATTAC-3’) and Rpl49-cDNA-R (5’-CTTGCGCTTCTTGGAGGAGA-3’), were used to synthesize cDNA contains ETS-18S, and Rpl49, respectively, from total RNA using SuperScript III Reverse Transcriptase (Invitrogen, #18080044). The cDNA product was diluted empirically to optimize amplification efficiency, melting curve, and appropriate Ct values. Diluted cDNA products were subjected to quantitative real-time PCR using SsoAdvanced^TM^ Universial SYBR^®^ Green Supermix (Bio-Rad) on a StepOnePlus real-time PCR system (Applied Biosystems). Relative rDNA transcript levels were calculated using the ^ΔΔ^Ct method and normalized to *RpL*32(*rp*49) mRNA levels. Primers used for RT-qPCR analysis of ETS and standard gene Rp49 are list in **Supplementary Table 1**.

### Immunofluorescence and Combined Immuno-FISH

#### Immunostaining

Embryos were collected on apple juice-agar plates and aged till the appropriate stage, dechorionated in 50% bleach, fixed in 1:1 heptane: 4% formaldehyde in 1×PBS for 25 mins, devitellinized in a 1:1 mixture of methanol: heptane, and stored at −20°C in methanol. Embryos were rehydrated by washing in 1×PBS containing 0.2% Triton (PBST), blocked for ∼1 hr with 2% BSA in PBST, then incubated with the primary antibody at 4°C overnight. Following washes with PBST, embryos were incubated with the appropriate secondary antibody at room temperature for 2 hrs, stained in DAPI, and mounted onto a slide using VectaShield (Vector Laboratories) mounting medium. All primary antibodies were used at 1:250 dilution and secondaries at 1:1 000. Imaging was performed on a Zeiss LSM880 Airyscan confocal microscopy (Airyscan Fast mode) with 63× oil immersion objective at room temperature.

#### Fluorescence in situ Hybridization

Following dechorionation, fixation, and devitellinization as described above, embryos were washed in 2×SSC containing 0.1% Tween-20 (2×SSCT) with increasing formamide concentrations (20%, 40%, then 50%) for 15 min each. 100 ng of DNA probes in 40 μl of hybridization solution (50% formamide, 3×SSCT, 10% dextran sulfate) was added, denatured together with the embryos at 95°C for 5 min and incubated overnight at 37°C. Following hybridization, embryos were washed twice in 2×SSCT for 30 mins at 37°C and thrice in PBST for 5 mins at room temperature. After completing washes, embryos were stained in DAPI and mounted onto a slide using VectaShield (Vector Laboratories) mounting medium.

#### Combined Immuno-FISH

For combined in situ detection of proteins and DNA sequences, immunofluorescence was performed first, embryos were post-fixed in 4% formaldehyde for 25mins, then processed for FISH.

### Microscopy

Imaging was performed on a Zeiss LSM 880 confocal microscopy equipped with an Airyscan detector, using Airyscan Fast mode and a 63× oil immersion objective (NA 1.4) at room temperature. Imaging parameters were kept constant across genotypes and experimental conditions within each experiment. Fluorophores were excited using 405 nm, 488 nm, 561 nm, and 633 nm laser lines. Images were acquired using Zeiss Zen Black software. Airyscan processing was performed using default settings in Zen, and reconstructed images were used for all subsequent analyses. Maximum intensity projections were generated from z stacks for visualization.

### Quantitative Image Analysis

All image analysis was performed on z-stack images acquired under identical imaging conditions across genotypes within each experiment, using Arivis Vision 4D (Zeiss) for segmentation and quantification unless specified otherwise.

#### H3K9me2 intensity

H3K9me2 levels were quantified in two ways. First, individual nuclei were segmented in 3D based on DAPI signal, and mean H3K9me2 fluorescence intensity was measured within each nucleus. Second, H3K9me2 intensity was quantified at the whole-embryo level. z-stacks were converted to maximum-intensity projections in ImageJ (Fiji), with each projected micrograph representing a single embryo containing multiple nuclei. Mean fluorescence intensity was measured across the full projected image using the ImageJ Measure tool, and the minimum pixel intensity within that same image was subtracted as a background estimate, yielding a single background-subtracted mean intensity value per embryo. Per-nucleus H3K9me2 intensity values appeared elevated in the Orc1 mutants relative to control embryos which was attributable to higher background fluorescence rather than increased signal. We therefore quantified this dataset using the background-subtracted whole-embryo method described above.

#### Nucleolar volume

Nucleoli were auto-segmented in 3D based on Fibrillarin or Modulo signal. Nucleolar volume was calculated from the resulting segmented objects.

#### rDNA-nucleolus overlap ratio

rDNA and nucleoli were independently segmented in 3D. Segmented rDNA objects were assigned to their corresponding nucleolus using a parent-child relationship in Arivis Vision 4D, and the intersection volume between each rDNA segment and its paired nucleolar segment was calculated. The overlap ratio was defined as the intersection volume divided by the total rDNA segment volume.

## Statistical Analysis

Quantitative image analysis measurements (H3K9me2 intensity, nucleolar volume, and rDNA-nucleolus overlap ratio) were obtained at the level of individual nuclei or segmented objects, with multiple measurements collected per embryo. Since measurements from the same embryo were not statistically independent, genotype comparisons were performed using linear mixed-effects models, in which genotype was included as a fixed effect and embryo as a random effect: *Measurement ∼ Genotype + (1 | Embryo).* Statistical analyses were performed in R using linear mixed-effects models implemented in the lme4 and lmerTest packages ^65^. This approach accounts for the non-independence among nuclei within the same embryo, while still retaining nucleus-level resolution rather than collapsing each embryo to a single average value. Pairwise comparisons between genotypes were performed using the emmeans package in R, and one-sided p values were calculated using the Satterthwaite approximation for degrees of freedom. For datasets consisting of bulk measurements rather than single-nucleus measurements, statistical significance was measured using a two-tailed Student’s *t*-test.

## Key resources table

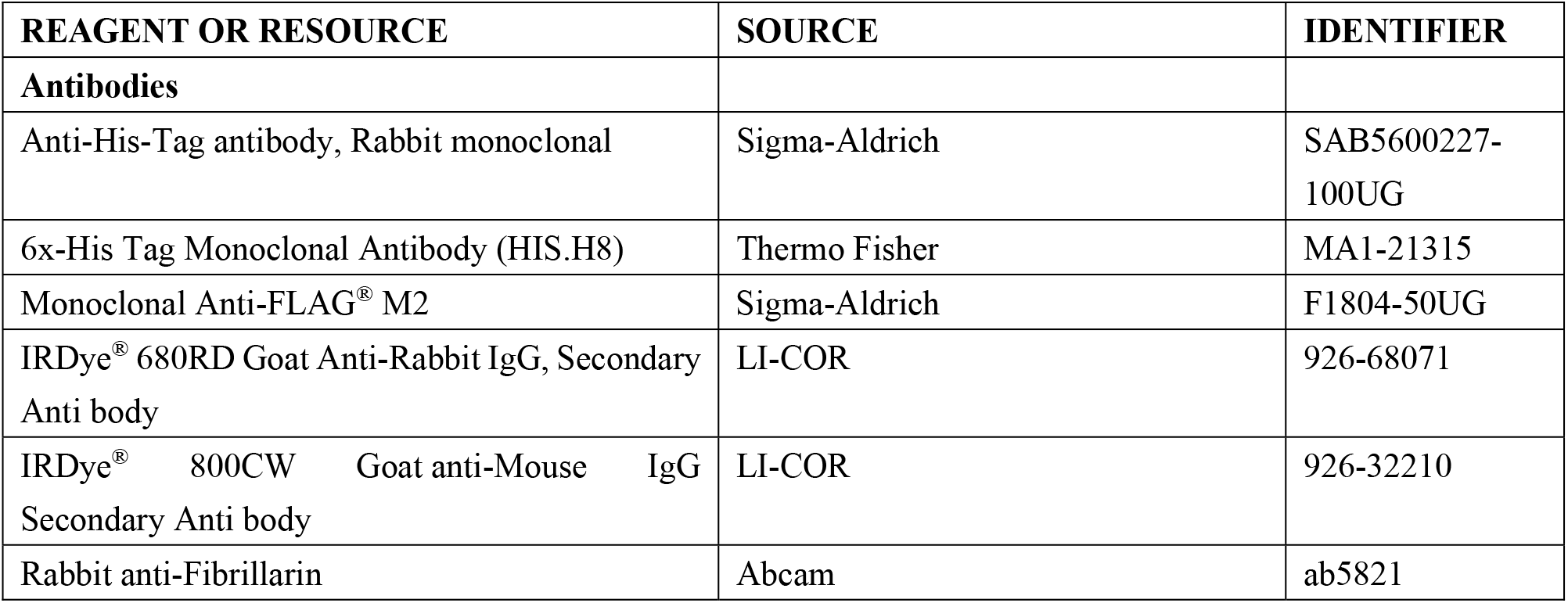
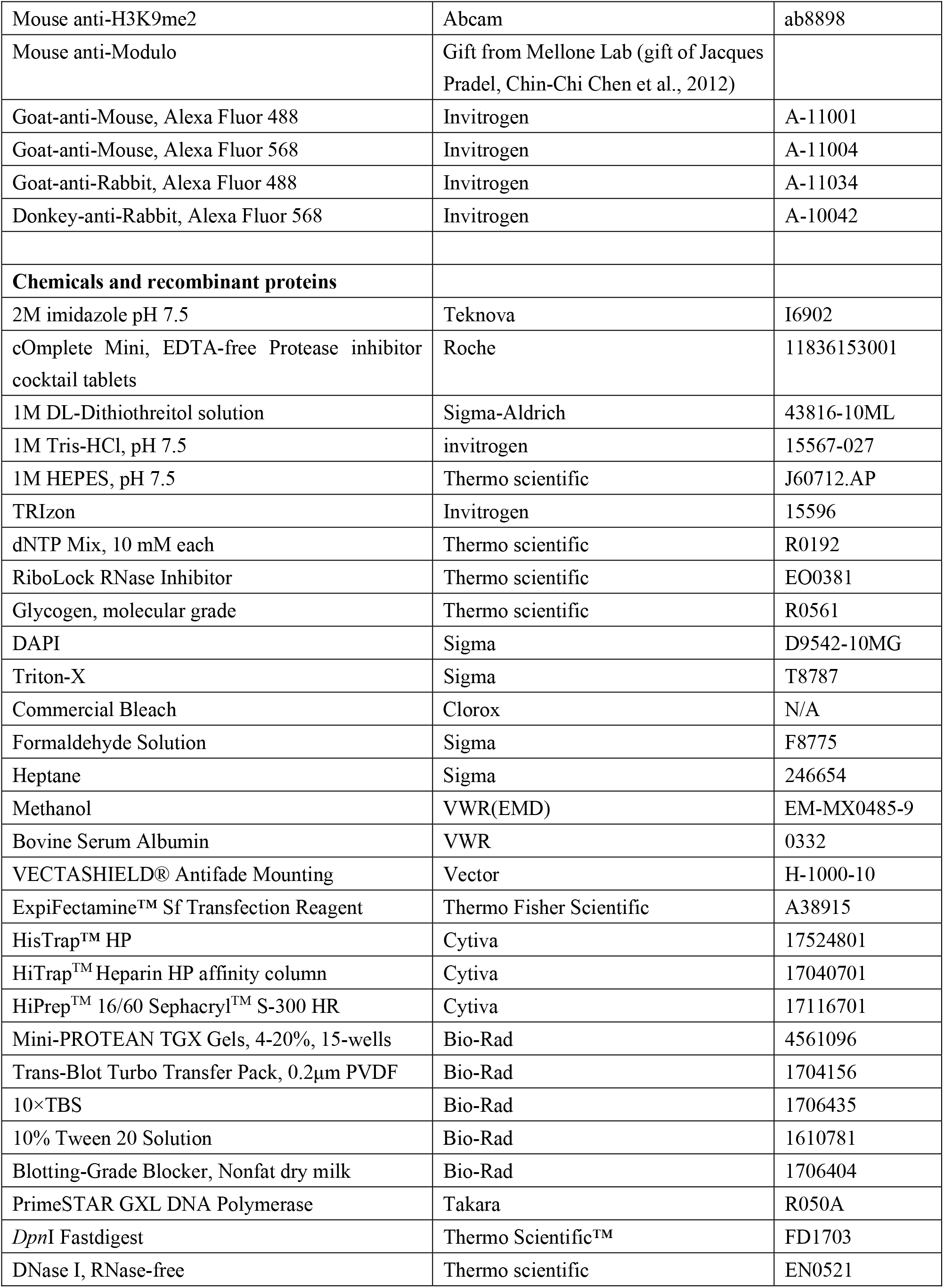
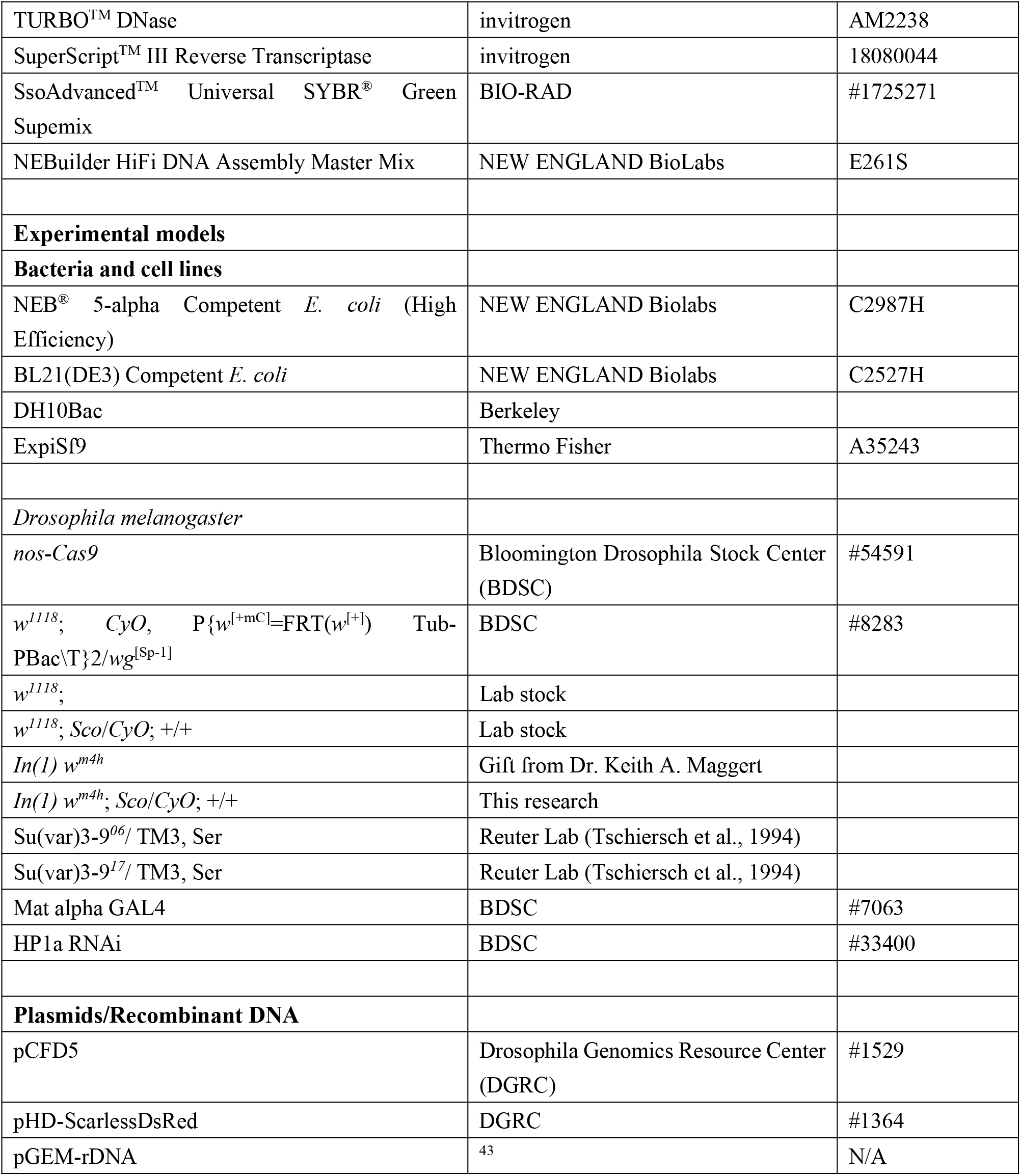

| REAGENT OR RESOURCE | SOURCE | IDENTIFIER |
| --- | --- | --- |
| **Antibodies** |  |  |
| Anti-His-Tag antibody, Rabbit monoclonal | Sigma-Aldrich | SAB5600227-100UG |
| 6x-His Tag Monoclonal Antibody (HIS.H8) | Thermo Fisher | MA1-21315 |
| Monoclonal Anti-FLAG® M2 | Sigma-Aldrich | F1804-50UG |
| IRDye® 680RD Goat Anti-Rabbit IgG, Secondary Anti body | LI-COR | 926-68071 |
| IRDye® 800CW Goat anti-Mouse IgG Secondary Anti body | LI-COR | 926-32210 |
| Rabbit anti-Fibrillarin | Abcam | ab5821 |
| Mouse anti-H3K9me2 | Abcam | ab8898 |
| Mouse anti-Modulo | Gift from Mellone Lab (gift of Jacques Pradel, Chin-Chi Chen et al., 2012) |  |
| Goat-anti-Mouse, Alexa Fluor 488 | Invitrogen | A-11001 |
| Goat-anti-Mouse, Alexa Fluor 568 | Invitrogen | A-11004 |
| Goat-anti-Rabbit, Alexa Fluor 488 | Invitrogen | A-11034 |
| Donkey-anti-Rabbit, Alexa Fluor 568 | Invitrogen | A-10042 |
| **Chemicals and recombinant proteins** |  |  |
| 2M imidazole pH 7.5 | Teknova | I6902 |
| cOmplete Mini, EDTA-free Protease inhibitor cocktail tablets | Roche | 11836153001 |
| 1M DL-Dithiothreitol solution | Sigma-Aldrich | 43816-10ML |
| 1M Tris-HCl, pH 7.5 | invitrogen | 15567-027 |
| 1M HEPES, pH 7.5 | Thermo scientific | J60712.AP |
| TRIzol | Invitrogen | 15596 |
| dNTP Mix, 10 mM each | Thermo scientific | R0192 |
| RiboLock RNase Inhibitor | Thermo scientific | EO0381 |
| Glycogen, molecular grade | Thermo scientific | R0561 |
| DAPI | Sigma | D9542-10MG |
| Triton-X | Sigma | T8787 |
| Commercial Bleach | Clorox | N/A |
| Formaldehyde Solution | Sigma | F8775 |
| Heptane | Sigma | 246654 |
| Methanol | VWR(EMD) | EM-MX0485-9 |
| Bovine Serum Albumin | VWR | 0332 |
| VECTASHIELD® Antifade Mounting | Vector | H-1000-10 |
| ExpiFectamine™ Sf Transfection Reagent | Thermo Fisher Scientific | A38915 |
| HisTrap™ HP | Cytiva | 17524801 |
| HiTrap™ Heparin HP affinity column | Cytiva | 17040701 |
| HiPrep™ 16/60 Sephacryl™ S-300 HR | Cytiva | 17116701 |
| Mini-PROTEAN TGX Gels, 4-20%, 15-wells | Bio-Rad | 4561096 |
| Trans-Blot Turbo Transfer Pack, 0.2µm PVDF | Bio-Rad | 1704156 |
| 10×TBS | Bio-Rad | 1706435 |
| 10% Tween 20 Solution | Bio-Rad | 1610781 |
| Blotting-Grade Blocker, Nonfat dry milk | Bio-Rad | 1706404 |
| PrimeSTAR GXL DNA Polymerase | Takara | R050A |
| *DpnI* Fastdigest | Thermo Scientific™ | FD1703 |
| DNase I, RNase-free | Thermo scientific | EN0521 |
| TURBO™ DNase | invitrogen | AM2238 |
| SuperScript™ III Reverse Transcriptase | invitrogen | 18080044 |
| SsoAdvanced™ Universal SYBR® Green Supemix | BIO-RAD | #1725271 |
| NEBuilder HiFi DNA Assembly Master Mix | NEW ENGLAND BioLabs | E261S |
| **Experimental models** |  |  |
| **Bacteria and cell lines** |  |  |
| NEB® 5-alpha Competent *E. coli* (High Efficiency) | NEW ENGLAND Biolabs | C2987H |
| BL21(DE3) Competent *E. coli* | NEW ENGLAND Biolabs | C2527H |
| DH10Bac | Berkeley |  |
| ExpiSf9 | Thermo Fisher | A35243 |
| *Drosophila melanogaster* |  |  |
| *nos-Cas9* | Bloomington Drosophila Stock Center (BDSC) | #54591 |
| $w^{1118}; CyO, P\{w^{[+mC]}=FRT(w^{[+]})\} Tub-PBac\{T\}2/wg^{[Sp-1]}$ | BDSC | #8283 |
| $w^{1118};$ | Lab stock | |
| $w^{1118}; Sco/CyO; +/+$ | Lab stock | |
| $In(1) w^{m4h}$ | Gift from Dr. Keith A. Maggert | |
| $In(1) w^{m4h}; Sco/CyO; +/+$ | This research | |
| Su(var)3-9<sup>06</sup>/ TM3, Ser | Reuter Lab (Tschiersch et al., 1994) |  |
| Su(var)3-9<sup>17</sup>/ TM3, Ser | Reuter Lab (Tschiersch et al., 1994) |  |
| Mat alpha GAL4 | BDSC | #7063 |
| HP1a RNAi | BDSC | #33400 |
| **Plasmids/Recombinant DNA** |  |  |
| pCFD5 | Drosophila Genomics Resource Center (DGRC) | #1529 |
| pHD-ScarlessDsRed | DGRC | #1364 |
| pGEM-rDNA | 43 | N/A |

## Supplemental information

**Supplementary Figure 1.**
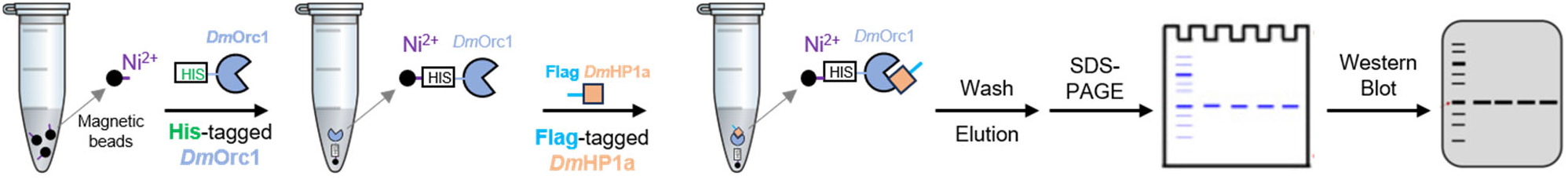
Schematic workflow of the Ni^2+^-bead pull-down assay used to detect binding between His-tagged *Dm*Orc1 fragments and FLAG-tagged *Dm*HP1a *in vitro*.

**Supplementary Figure 2.**
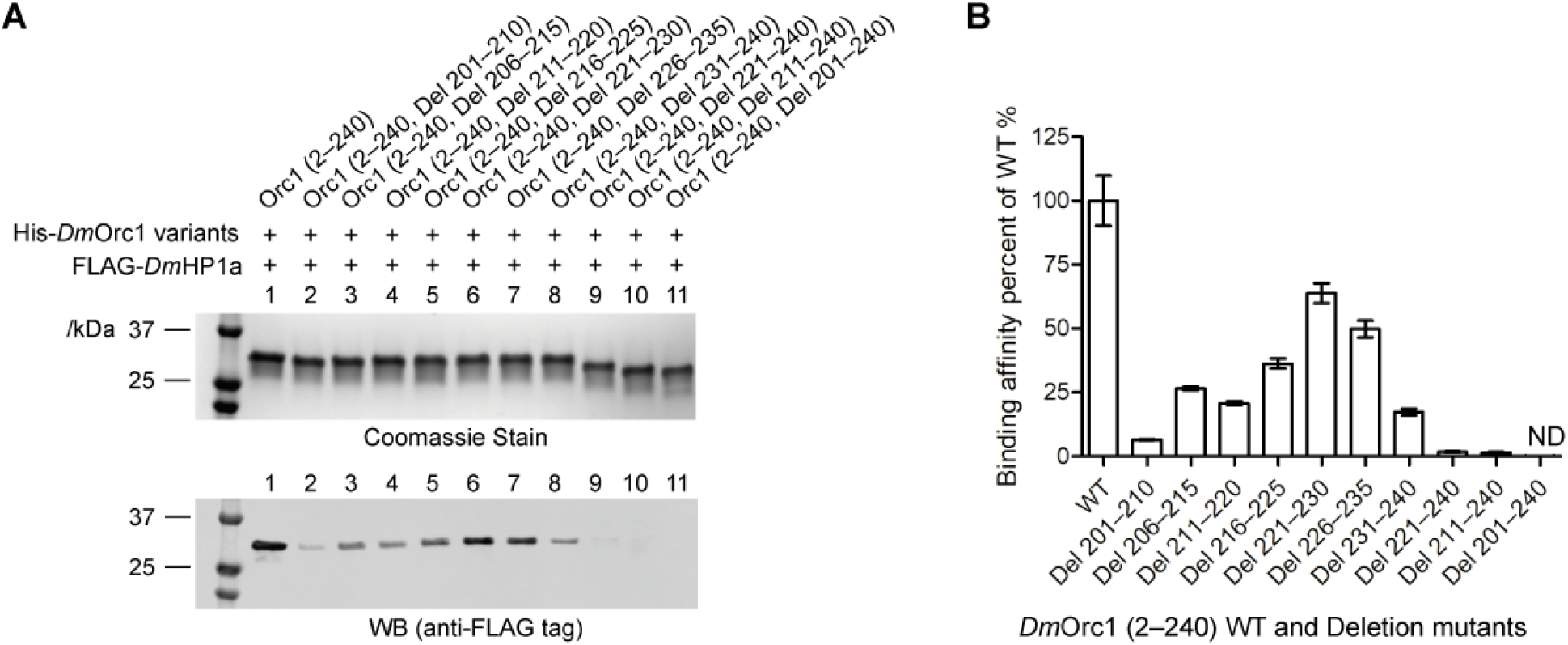
HP1a binding affinities of Orc1 variants generated by overlapping deletions or C-terminal truncations within the His-tagged Orc1 (2–240) fragment (**A**) and the quantification of binding affinity compared to wild-type (n=3, **B**). Deleted regions are indicated as defined in Figure 1C. ND=not detected.

**Supplementary Figure 3.**
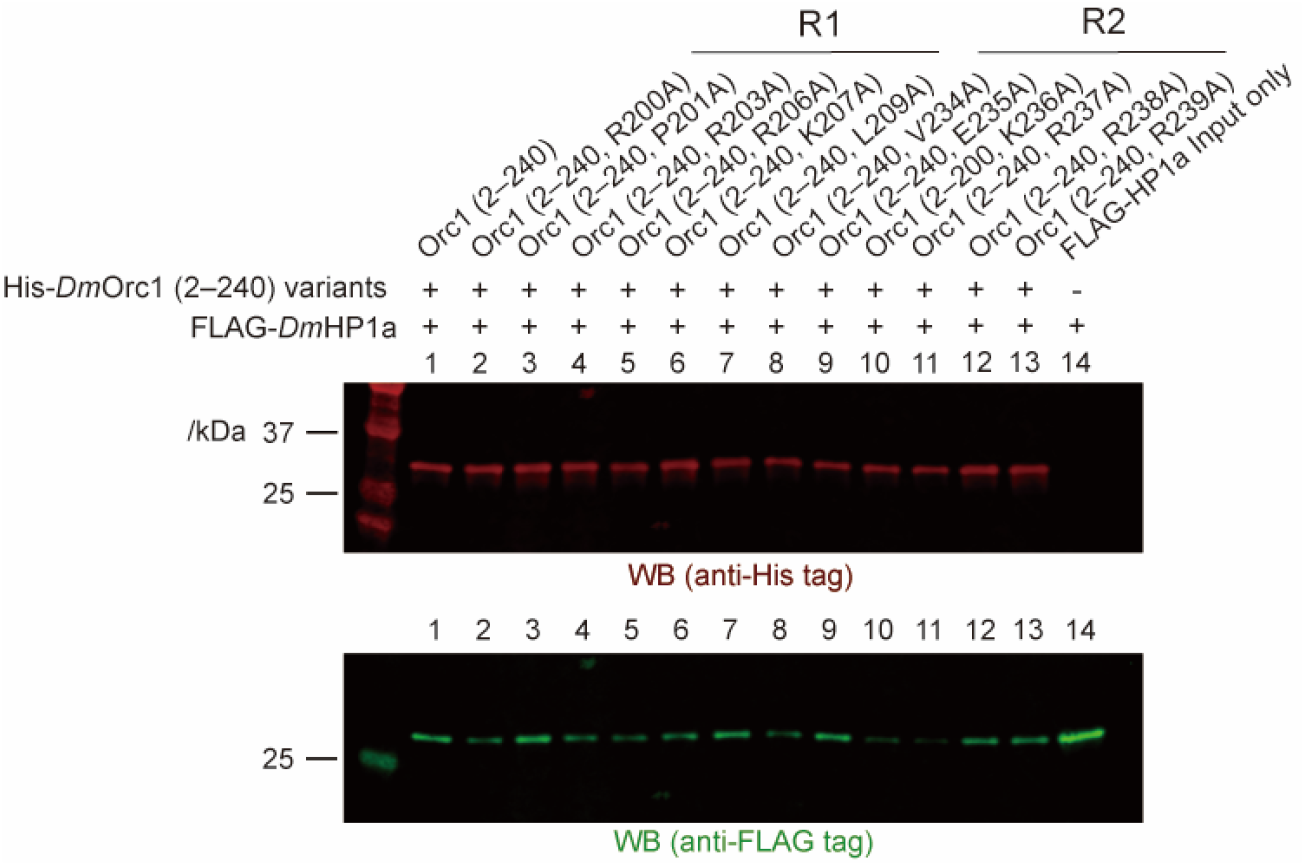
Representative binding results of single alanine-substituted Orc1 (2–240) variants to HP1a, relative to wild-type Orc1 (2–240). Alanine scanning targeted residues with bulky side chains within R1 (residues 200–209) and R2 (residues 234–239) in the *Dm*Orc1 IDR.

**Supplementary Figure 4.**
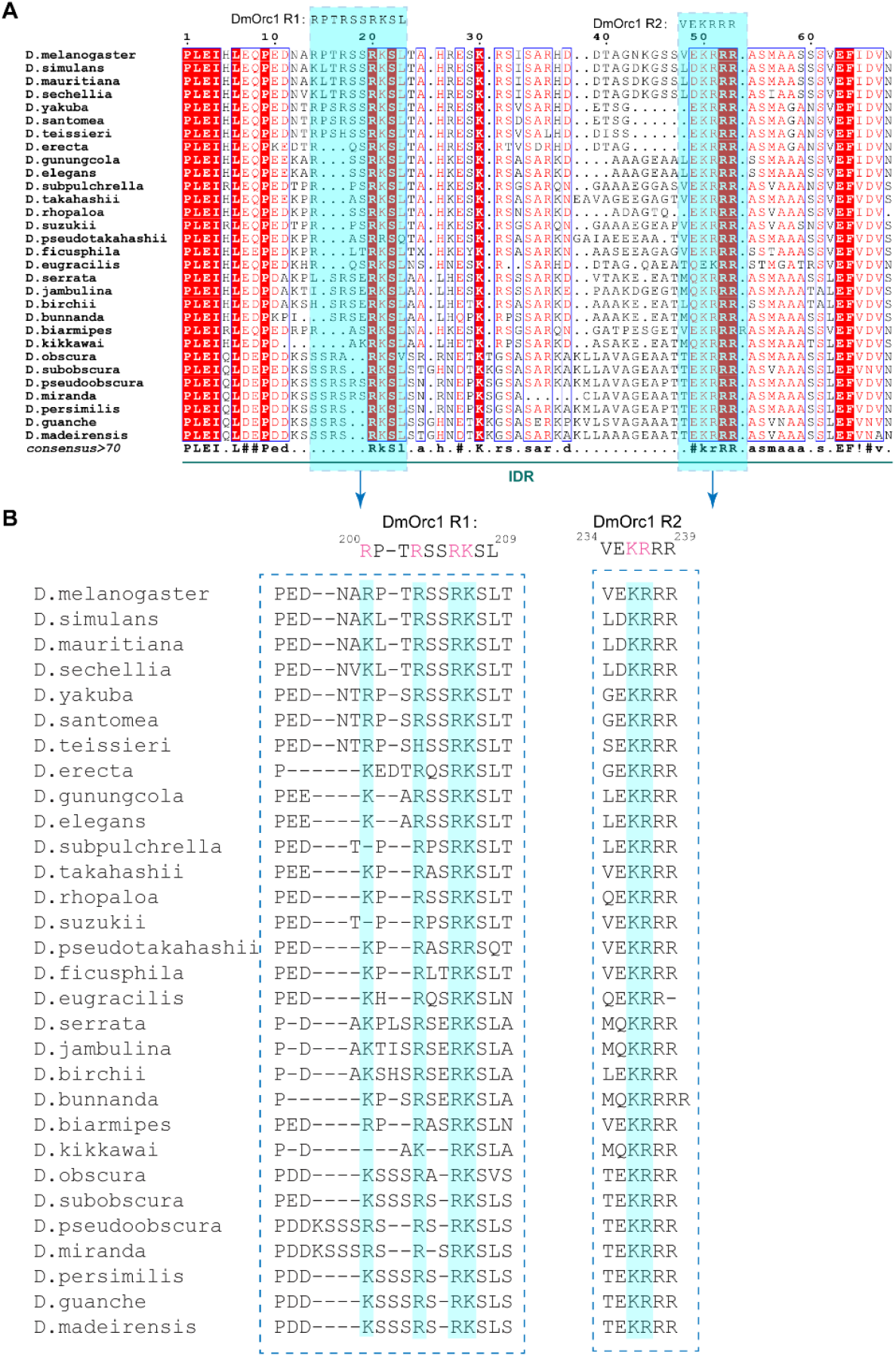
Sequence alignment of R1- and R2-containing regions within the Orc1 IDR across different *Drosophila* species. **(A)** Alignment of the Orc1 IDR segment containing R1 and R2 from multiple *Drosophila* species. **(B)** Alignment of the R1 and R2 motifs across *Drosophila* species. R/K residues in magenta on the top in R1 and KR in R2 of Orc1 are critical residues that involves in HP1a binding.

**Supplementary Figure 5.**
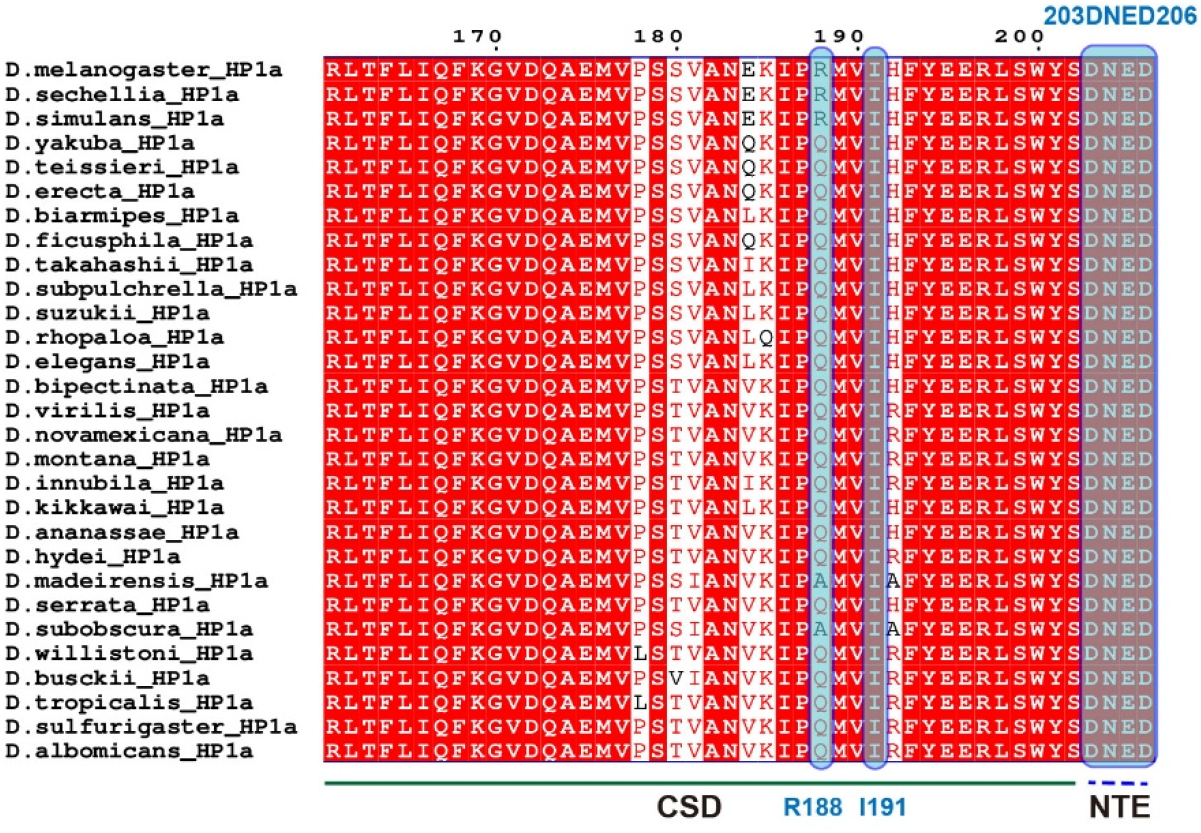
Alignment of the HP1a C-terminal region from *Drosophila* species, including part of chromoshadow domain (CSD) and the C-terminal extension (CTE). The *D. melanogaster* CTE, containing residues 203–206 (DNED), and residue I191, which regulates the monomer–dimer transition, are both highly conversed across *Drosophila*. Residue R188 is predominantly substituted with glutamine (Q) in most species (a change associated with enhanced dimerization), with alanine (A) observed in a minority of cases (n=2).

**Supplementary Figure 6.**
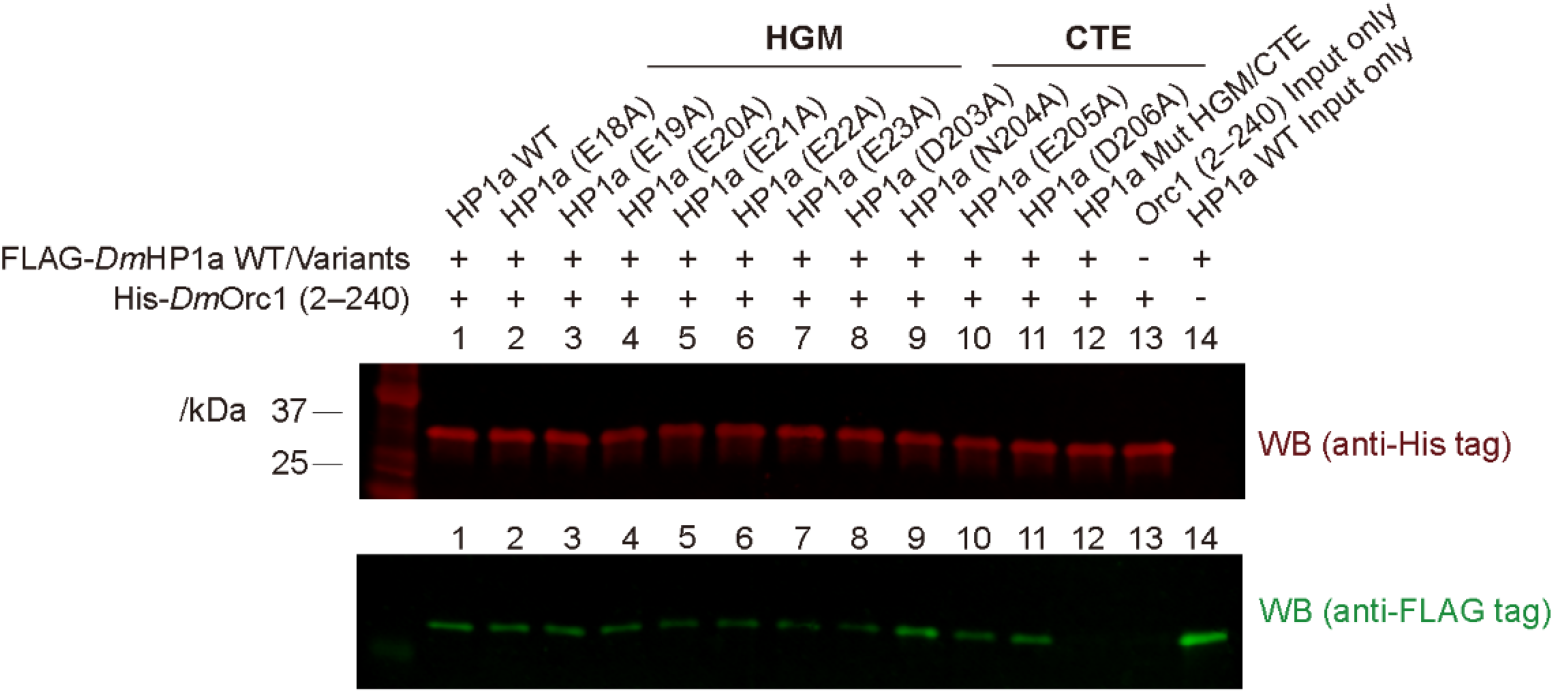
Representative binding results of single alanine-substituted HP1a variants to Orc1 (2–240), relative to wild-type HP1a. Alanine scanning targeted residues with bulky side chains within HGM (residues 18–23) and CTE (residues 203–206) in HP1a. The combined mutant HP1a (Mut HGM/CTE) represents all residues in HGM and CTE were mutated to alanine.

**Supplementary Figure 7.**
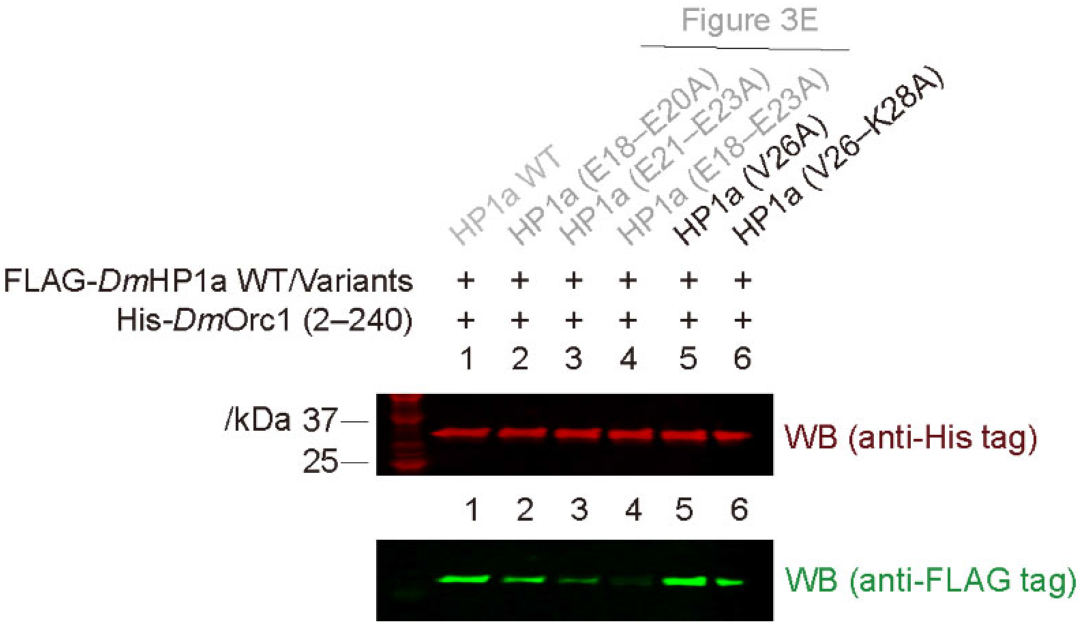
Binding results of HP1a variants V26A and V26–K28A (residues 26–28, VEK) to Orc1 (2–240). The Orc1 (2–240) binding results of HP1a variants (E18–E20A), (E21–E23A), and (E18–E23A) were shown in Figure 3E.

**Supplementary Figure 8.**
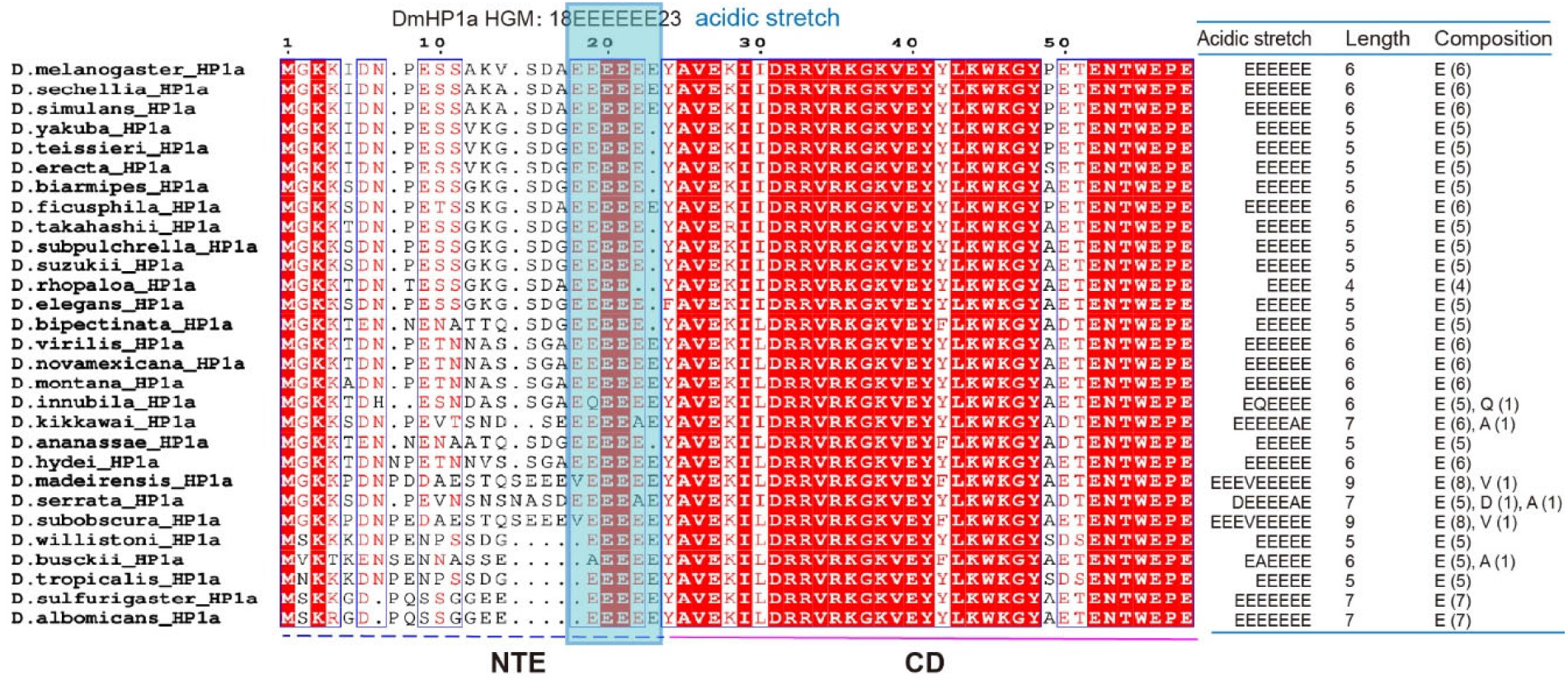
Alignment of the HP1a N-terminal region from *Drosophila* species, including the N-terminal extension (NTE) and part of the chromodomain (CD), revealing a conserved acidic stretch locates at the upstream of the CD that resembles the hexa-glutamic acid motif (HGM) in *Drosophila melanogaster*. The length distribution of acidic stretch is dominated by 5 aa (n=12, 41%) and 6 aa (n=10, 35%), with minor contributions from 7 aa (n=4, 14%), 9 aa (n=2, 14%), and 4 aa (n=1, 3%). The sequences are mostly composed of glutamic acid (E, n=163, 96%), with only minor contribution of alanine (A, n=3), valine (V, n=2), aspartic acid (D, n=1), and glutamine (Q, n=1), highlighting a strongly biased E composition.

**Supplementary Figure 9.**
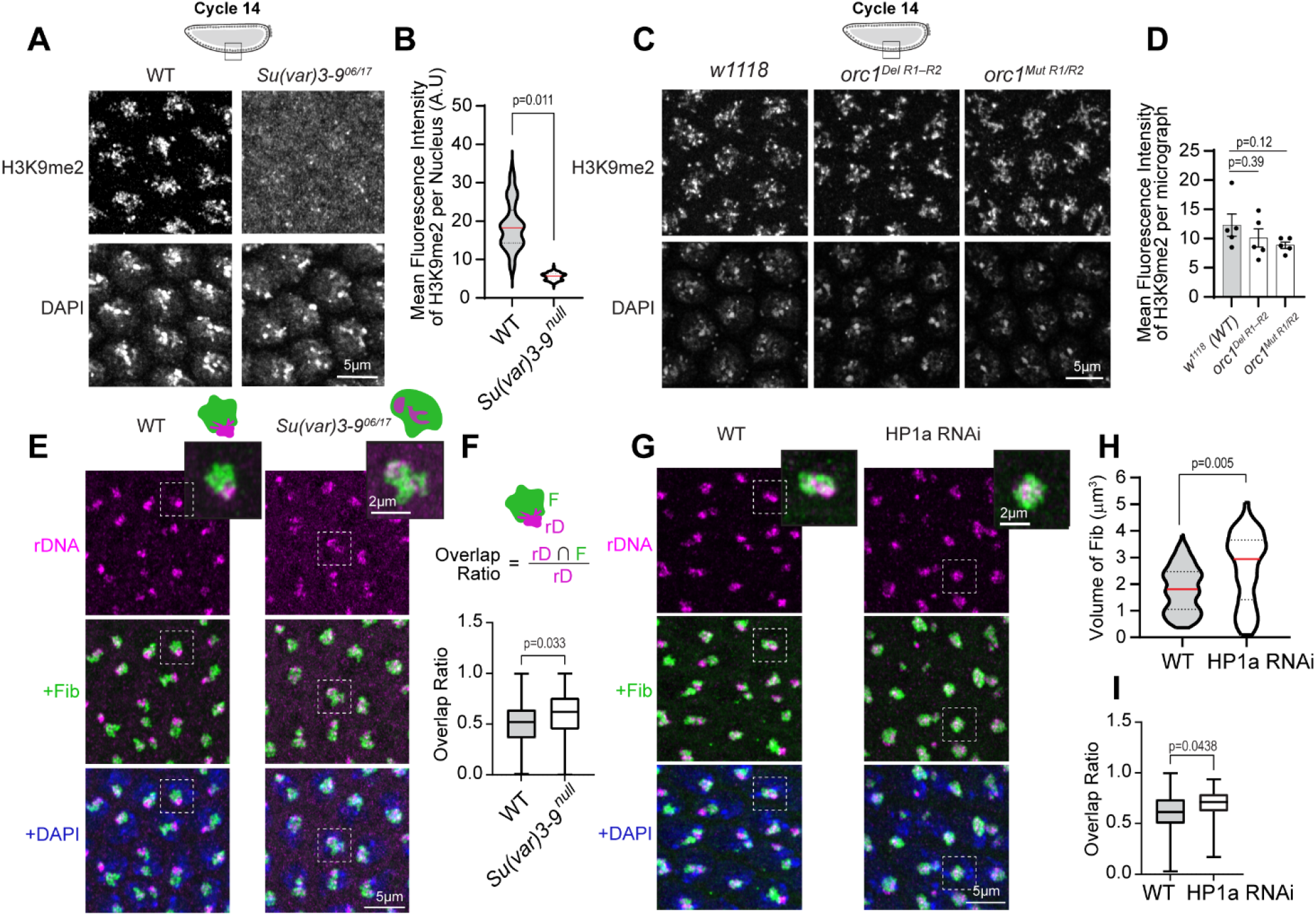
Reduced H3K9 methylation or HP1a levels lead to rDNA decondensation and nucleolar enlargement. **(A)** Maximum intensity z projections of nuclei from wild-type (WT, Oregon R) and *Su(var)3-9*^null^ embryos immunostained for H3K9me2 and counterstained with DAPI (blue) at nuclear cycle 14. *Su(var)3-9*^null^ embryos show a reduction in H3K9me2 signal relative to WT. **(B)** Violin plot indicating the distribution of mean intensities of H3K9me2 fluorescence per nucleus in WT and *Su(var)3-9*^null^ embryos at nuclear cycle 14, with width indicating density of data points. Horizontal lines indicate median (red solid line) and quartiles (black dashed lines). *OreR/WT*, n = 210 nuclei from 4 embryos; *Su(var)3-9* ^null^, n = 137 nuclei from 3 embryos. Statistical comparisons were performed using a linear mixed-effects model with genotype as a fixed effect and embryo as a random effect, to account for non-independence of nucleoli within the same embryo (See Methods for details). **(C)** Maximum intensity z projections of nuclei from *w*^1118^ and *Orc1* mutant embryos stained for H3K9me2 and DAPI at nuclear cycle 14. H3K9me2 signal intensity and distribution appear comparable between genotypes. **(D)** Quantification of mean H3K9me2 fluorescence intensity per micrograph in *w*^1118^/WT, *orc1^Del^ ^R1–R2^*, and *orc1^Mut^ ^R1/R2^* embryos. Each point represents a micrograph from one embryo with n = 5 embryos for each genotype. Bars indicate mean ± SEM and p values were calculated using two-tailed Student’s t-tests. **(E)** Combined immuno-FISH maximum intensity z projections of embryos stained for the nucleolar marker Fibrillarin (Fib,green), rDNA (magenta), and DNA (DAPI, blue) in WT and *Su(var)3-9*^null^ embryos. Loss of H3K9 methylation is associated with redistribution of rDNA and enlarged nucleolar morphology. **(F)** Quantification of the rDNA overlap ratio in WT and *Su(var)3-9* ^null^ embryos. The overlap ratio was calculated as the fraction of the rDNA (rD) signal overlapping the nucleolar (Fibrillarin, F) marker, as illustrated above. Box plots show the median, interquartile range, and minimum to maximum values. *OreR/WT*, n = 981 rDNA-nucleolar pairs from 3 embryos; *Su(var)3-9* ^null^, n = 1033 rDNA-nucleolar pairs from 3 embryos. P value was calculated using a linear mixed-effects model with genotype as a fixed effect and embryo as a random effect. **(G)** Maximum intensity z projections of nuclei from WT and HP1a RNAi embryos at nuclear cycle 14 stained for Fibrillarin (Fib, green), rDNA (magenta), and DNA (DAPI, blue). HP1a depletion phenocopies the rDNA decondensation and nucleolar enlargement observed in *Su(var)3-9*^null^ embryos. **(H)** Violin plots show the distribution of nucleolar volumes based on Fibrillarin staining in WT and HP1a RNAi embryos at nuclear cycle 14, with width indicating density of data points. Horizontal lines indicate median (red, solid) and quartiles (black, dashed). *OreR/WT*, n = 242 nucleoli from 5 embryos; *HP1a RNAi*, n = 180 nucleoli from 5 embryos. P value was calculated using a linear mixed-effects model. **(I)** Quantification of rDNA overlap ratio in WT and HP1a RNAi embryos, measured as in (F). *OreR/WT*, n = 244 rDNA-nucleolar pairs from 5 embryos; *HP1a RNAi*, n = 175 rDNA-nucleolar pairs from 5 embryos.

**Supplementary Figure 10.**
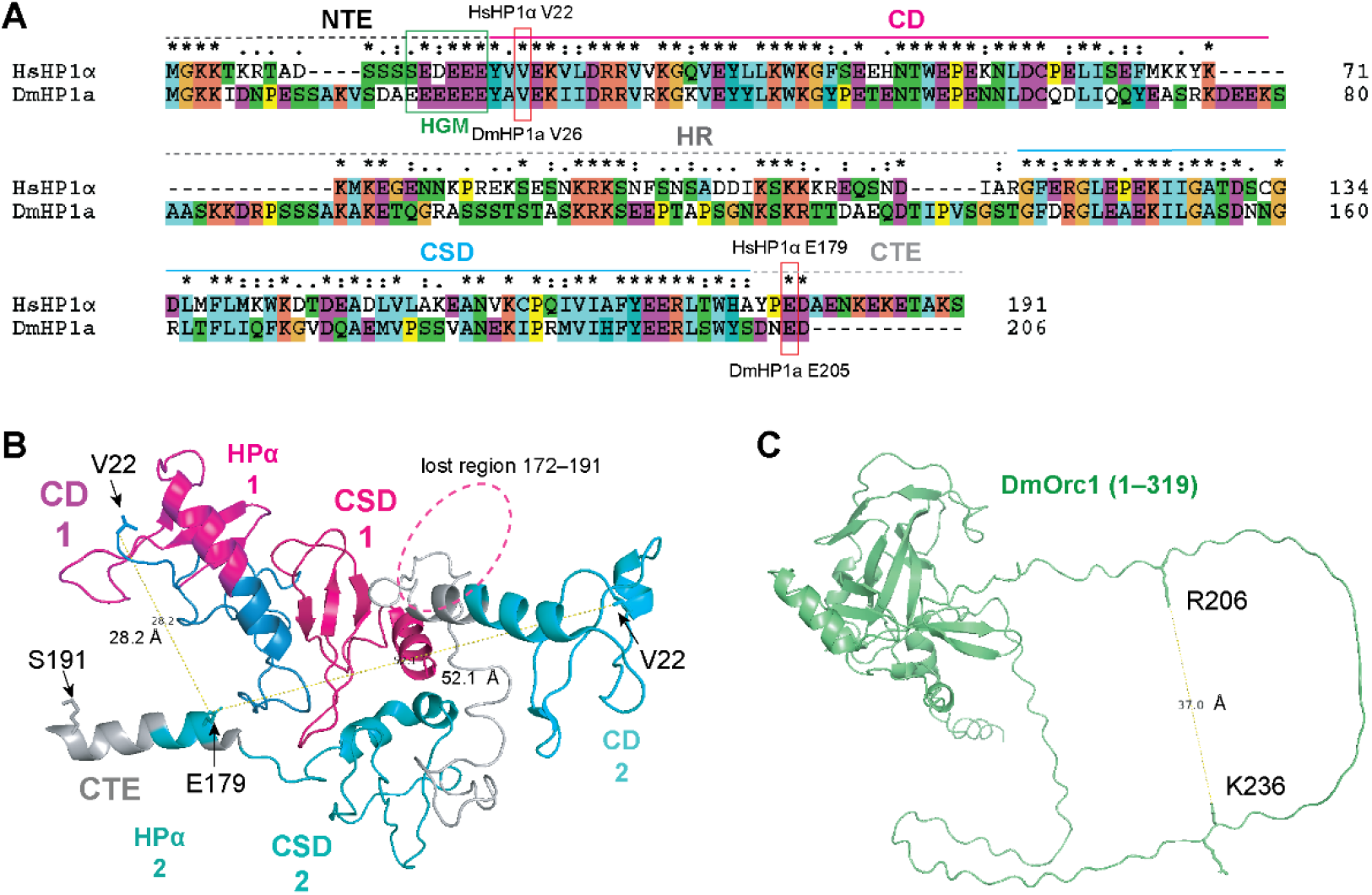
Structural comparison of cis and trans binding models for Orc1 recognition of the HP1 dimer. (A) Sequence alignment of human HP1α (HsHP1α) and *Drosophila* HP1a (DmHP1a) showing the correspondence between residues for distance measurements. HsHP1α V22 corresponds to DmHP1a V26, and HsHP1α E179 corresponds to DmHP1a E205. (B) Distance measurements based on cryo-EM structure of human HP1α dimer bound to an H2A.Z-nucleosome ^47^ [1] (PDB ID: 8UXQ). The two HP1α protomers are shown in distinct colors: HP1α1 (CD1 in purple and CSD1 in hot pink) and HP1α2 (CD2 in cyan and CSD2 in teal). Residues 1–21 of the N-terminal extension (NTE) are unresolved in both protomers. In HP1α1, residues 172–191 are unresolved and indicated as a lost region. In HP1α2, the C-terminal extension (CTE) is shown in gray, with residues E179–E182 (EDAE, in cyan) within this region are homologous to the *Drosophila* HP1a CTE (203–206, DNED). Because the conserved HGM in human HP1α is located within the unresolved N-terminal extension, V22, the first resolved residue adjacent to the HGM, was used as a surrogate for distance measurements; HsHP1α E179 locates in CTE and was chosen. Distances were measured from V22 of both HP1α protomer to E179 of HP1α 2 subunit. The distances between V22 and E179 within HP1α2 represent the cis-binding configuration, whereas the distance between V22 of HP1α1 and E179 of HP1α2 represents trans-binding configuration. (C) The distance between R206 (R1) and K236 (R2) in the AlphaFold-predicted structure of DmOrc1 (1–319).

---

**Supplementary Table 1.**
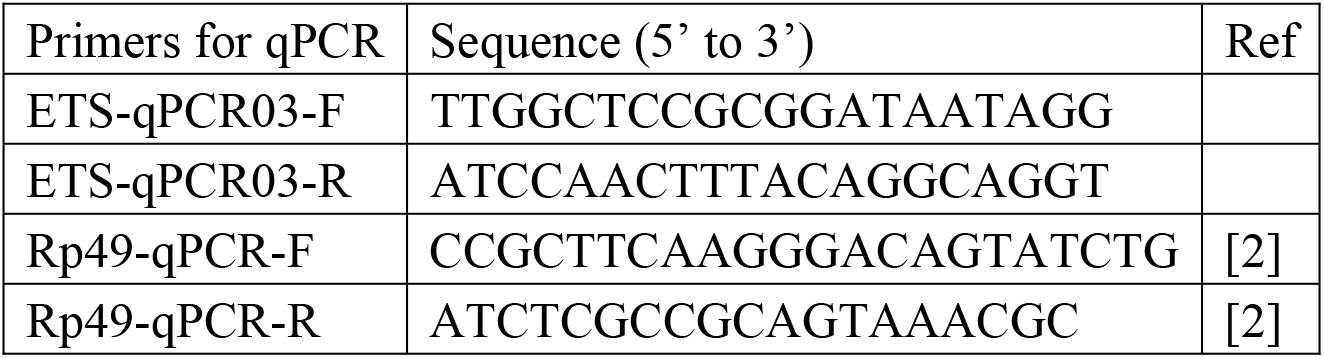
Primers used to RT-qPCR.

